# Identification of Mebendazole as an ethylene signaling activator reveals a role of ethylene signaling in the regulation of lateral root angles

**DOI:** 10.1101/2022.02.16.480742

**Authors:** Wenrong He, Hai An Truong, Ling Zhang, Min Cao, Neal Arakawa, Yao Xiao, Kaizhen Zhong, Yingnan Hou, Wolfgang Busch

## Abstract

The lateral root angle or gravitropic set-point angle (GSA) is an important trait for root system architecture (RSA) that determines the radial expansion of the root system. The GSA therefore plays a crucial role for the ability of plants to access nutrients and water in the soil. Despite its importance, only few regulatory pathways and mechanisms that determine GSA are known, and these mostly relate to auxin and cytokinin pathways. Here, we report the identification of a small molecule, Mebendazole (MBZ) that modulates GSA in *Arabidopsis thaliana* roots and acts via the activation of ethylene signaling. We uncover that MBZ directly acts on the serine/threonine protein kinase CTR1, which is a negative regulator of ethylene signaling. Our study not only reveals that the ethylene signaling pathway is essential for GSA regulation, but it also identifies a small molecular modulator of RSA and the first small molecule that acts downstream of ethylene receptors and that directly activates ethylene signaling.

**In brief:** He *et al*. identify a small molecule that regulates lateral root angle. They show that the compound increases lateral root angle by inhibiting CTR1 kinase activity, which in turn activates ethylene signaling. Therefore, they uncover that the ethylene pathway is involved in lateral root angle regulation.

**Highlights:**

- MBZ increases lateral root angle
- MBZ regulates lateral root angle by activating ethylene signaling
- MBZ inhibits CTR1 kinase activity
- The ethylene pathway is involved in lateral root angle regulation

## INTRODUCTION

The root systems of plants are vital for plant survival and productivity. They anchor plants and they absorb water and nutrients from the soil. The distribution of roots in the soil, root system architecture (RSA), is of enormous relevance. For instance, deeper rooting is associated with higher carbon permanence (Poirier et al., 2018), a critical trait for soil carbon sequestration and the ability of plants to survive terminal droughts (Uga et al., 2013). Besides, to enhance the capacity for root systems to efficiently uptake nitrogen and water, deeper roots are required, yet for improving the uptake of phosphate and other nutrients, shallower soil layers have to be accessed by roots (Lynch, 2019). A key determinant of RSA is the lateral root angle or gravitropic set-point angle (GSA). Previous studies have shown that auxin plays a crucial role in regulating GSA (Rosquete et al., 2013a; Roychoudhry et al., 2013; Xiao and Zhang, 2020). Application of exogenous IAA or synthetic auxin leads to lateral root growth toward the vector of gravity thereby decreasing GSA (Rosquete et al., 2013a; Roychoudhry et al., 2013). Another phytohormone, cytokinin, was also found to be involved in the regulation of GSA. Application of cytokinin showed an anti-gravitropic effect and thereby increased GSA (Waidmann et al., 2019b). Finally, members of the IGT gene family, including DRO1/LAZY4 have been shown to play a major role in lateral root angles via auxin associated mechanisms (Ge and Chen, 2016; Guseman et al., 2017; Waite and Dardick, 2021; Yoshihara and Spalding, 2017).

Ethylene is an ancient plant hormone. The ethylene signaling pathway is highly conserved and was present already in the aquatic progenitors of land plants (Ju et al., 2015). Ethylene is produced from the conversion of S-adenosyl-L-methionine (SAM) to 1- aminocy-clopropane-1-carboxylic acid (ACC) and then to ethylene, which is catalyzed by enzymes including ACC synthase (ACS) and ACC oxidase (ACO) (Pattyn et al., 2021; S F Yang and Hoffman, 1984). When ethylene binds to its receptors (ETRs) (Chang et al., 1993; Hua et al., 1998; Sakai et al., 1998), it inactivates a serine/threonine protein kinase, CONSTITUTIVE TRIPLE RESPONSE1(CTR1) (Kieber et al., 1993). In the absence of ethylene, CTR1 blocks ethylene downstream responses by phosphorylating the C-terminal end of ETHYLENE-INSENSITIVE2 (EIN2), an ER–associated membrane protein that works as a positive regulator of ethylene signaling (Alonso et al., 1999; Ju et al., 2012; Wen et al., 2012). This leads to the degradation of EIN2-CEND by the Ub/26S proteasome (Ju et al., 2012; Qiao et al., 2012a; Wen et al., 2012). When CTR1 is inactivated by ethylene binding to ETRs, EIN2-CEND is stabilized and moves into the nucleus to transduce the ethylene signal to downstream transcription factors, such as EIN3, EIN3 LIKE1 (EIL1) (An et al., 2010; Chao et al., 1997) and ETHYLENE RESPONSE FACTORs (ERFs) (Muller and Munne-Bosch, 2015). These transcription factors activate the downstream ethylene response. The ethylene pathway was found to play an important role in numerous growth and developmental processes in the root, such as the inhibition of root elongation (Kieber et al., 1993; Le et al., 2001; Vaseva et al., 2018), induction of root hair growth (Feng et al., 2017), and inhibition of the initiation of lateral roots (LRs) (Negi et al., 2008).

To identify potential regulators of GSA, we performed a chemical genetics screen and identified a small molecule, Mebendazole (MBZ), which dramatically increased GSA. Genetic analysis revealed that MBZ increased GSA through rapidly activating the ethylene signaling pathway. Both MBZ and ACC treatment induced ethylene signaling, which led to a modulation of lateral root angle. Further molecular and biochemistry studies showed that MBZ activates ethylene signaling by specifically targeting CTR1 and thereby blocking its kinase activity. Our study not only identified MBZ as an efficient small molecular activator of ethylene signaling, but also uncovered that ethylene signaling plays an important role in regulating GSA.

## Results

### A small compound profoundly affects the lateral root gravitropic set-point angle (GSA) in Arabidopsis

To identify small molecules that affect GSA, we screened chemicals for their ability to change GSA of young seedlings from a small molecular library with 2000 diverse generic chemicals (SP 2000)(He et al., 2011; Sun et al., 2017; Zhu et al., 2019). We found that treatment with Mebendazole (MBZ) (Figure S1A) led to a distinct increase of lateral root angles that caused lateral roots to grow in a more horizontal direction (Figure 1A). The lateral root GSA of untreated two-week-old Arabidopsis seedlings was approx. 46°, and the GSA of lateral roots of seedlings grown on 1 µM MBZ plates was approx. 70° (Figure 1A, B). While the lateral root angle was the most striking phenotype, at a much earlier developmental stage (one week after germination), seedlings on 1 µM MBZ displayed a decreased root length (in average reduced by 37%) and an increased root hair initiation and elongation (Figure 1C, D, Figure S1B, C). To test how quickly MBZ would affect lateral root growth, we grew seedlings for 14 days on ½ MS medium with DMSO, then transferred them to ½ MS medium supplemented with 1 µM MBZ and acquired images every 10 minutes. All of the 5 measured lateral root tips started to grow at a more horizontal (less steep) angle within 3 hours of MBZ treatment (Figure 1E upper panel, F), which coincided with the first timepoint at which root angles were significantly different between control and MBZ (Welch Two Sample t-test; p-value <0.05). To test whether the action of MBZ was reversible, we germinated and grew seedlings on ½ MS for 5 days, and then transferred them to medium containing 1 µM MBZ for 14 days, thereby inducing lateral roots of MBZ treated seedlings to grow more horizontally. We note that we used this later timepoint (compared to the data presented in the upper panel) to be able to consider enough lateral roots that were growing at the more horizontal angle. We then transferred the seedlings back to control conditions and acquired images every 10 minutes. Already after one hour root angles were significantly different between control (MBZ) and treatment (removal from MBZ) conditions (Welch Two Sample t-test; p-value <0.05), and within 4 hours all 5 lateral root tips we measured started to grow at a distinctly steeper angle upon removal from MBZ (Figure 1E bottom panel, F). These data showed that the treatment of Arabidopsis roots with MBZ changes lateral root angles within 3 hours and that this effect is fully reversible. In summary, we found that MBZ changes lateral root angles and important RSA traits associated with the development of a shallow root system.

**Figure 1.**
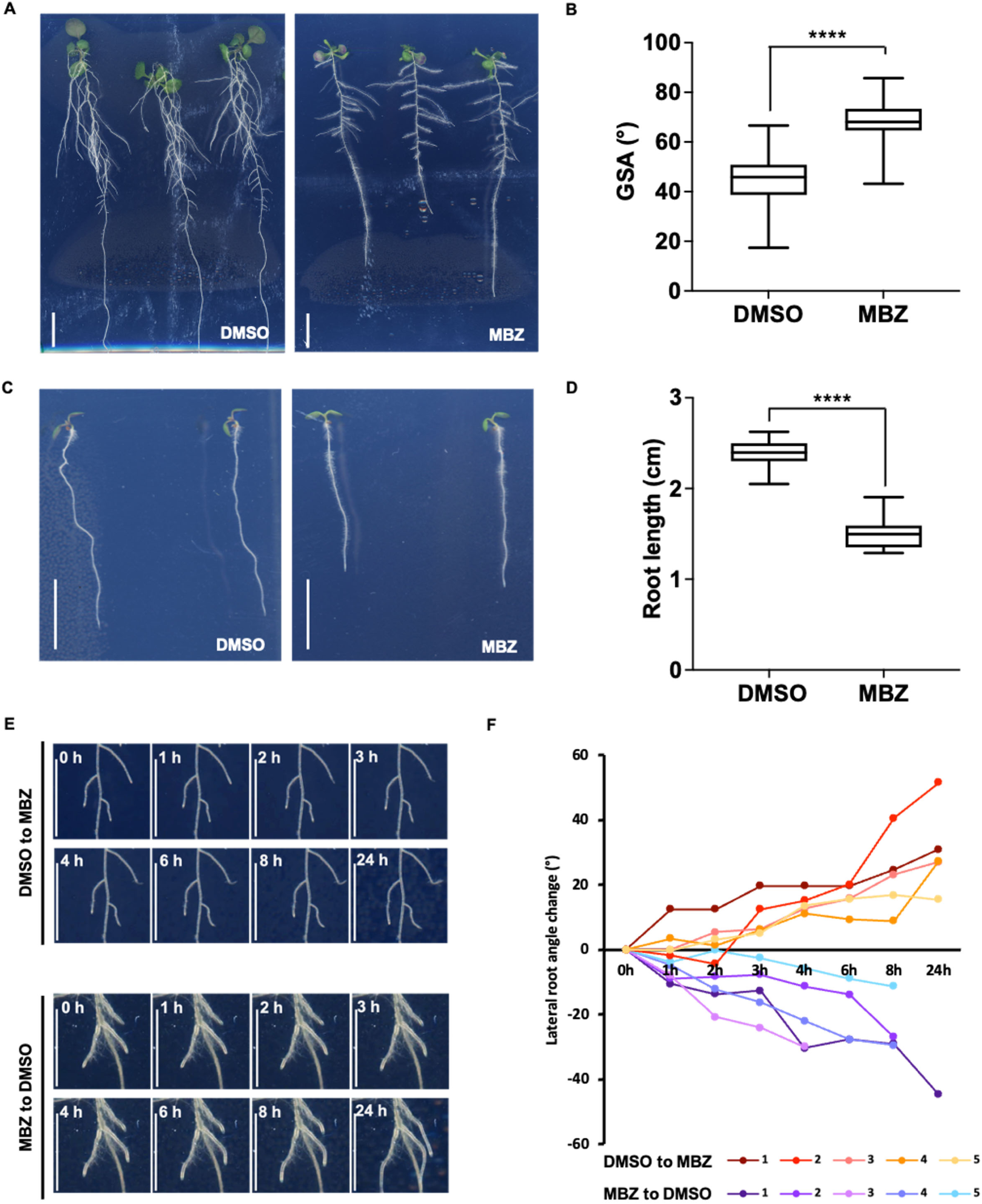
MBZ treatment perturbs root system architecture (RSA). (A) 14-day-old Arabidopsis seedlings of grown on DMSO and MBZ (1 µM) plates. (B) Quantification of GSA in (A). 15 LRs per seedling on DMSO plates, 12 LRs per seedling on MBZ plates, and 6 seedlings for each condition were used. Unpaired, two-tailed Student’s t-tests was used for statistical analysis. **** p < 0.0001. (C) 7-day-old seedlings on DMSO and MBZ (1 µM) plates. (D) Quantification of primary root length (cm) in (B). 15-20 seedlings for each treatment were used. **** p < 0.0001. (E) 14-day-old (upper panel) Arabidopsis seedlings on DMSO plates were transferred to MBZ (1 µM) plates, and 19-day-old (bottom panel) seedlings on MBZ (1 µM) plates were transferred to DMSO plates, followed by continuous scanning for 24 h. (F) Changes of lateral root angle after plate transfer from DMSO to MBZ (red and yellow lines) or MBZ to DMSO (blue, purple lines) compared to timepoint 0. (Scale bar: A, C, G, 1 cm; E, 5 mm.). Boxplots: Whiskers: min/max values; hinges: 25^th^ to 75^th^ percentile; mid-line: median.

### Application of MBZ induces the ethylene pathway

To identify the pathway that MBZ acted upon, we utilized a genome-scale expression profiling approach. For this, we performed an mRNA-seq experiment using whole roots of 14-day-old Arabidopsis grown on ½ MS media treated with 10 µM MBZ for 4 hours. The analysis revealed that 677 genes were induced compared to mock treatment (FDR<0.05; log2 fold change > 1) and 658 genes were repressed (FDR<0.05; log2 fold change < -1). A gene ontology (GO) analysis revealed that among the genes that were upregulated by MBZ, genes involved in ethylene responses were strongly overrepresented in this early response. In particular, the GO process “negative regulation of ethylene-activated signaling pathway” (GO:0010105) was the most enriched process (∼21 fold enrichment, P-value: 6.07*10^-4^) upon MBZ treatment (Figure 2A), and the 4^th^ most enriched GO process was “ethylene-activated signaling pathway” (GO:0009873) (∼11 fold enrichment, P-value:7.45*10^-5^) (Figure 2A). Other GO categories were also significantly enriched, most notably responses to light. However, given that ethylene signaling was the most enriched GO term, we first looked at the responses of genes involved in the ethylene hormone pathway. A major portion of these genes were differentially regulated by MBZ compared to mock treatment (Figure 2A, B). We therefore hypothesized that MBZ elicited the ethylene pathway. To test this hypothesis, we first compared the set of genes that had responded to MBZ with a published transcriptome dataset that had been generated by exposing roots to 4 hours of ethylene treatment (Chang et al., 2013). Of the 1335 differentially expressed genes (DEGs) upon the 4 hours of MBZ treatment, 306 were overlapping with 1397 DEGs upon 4 hours of ethylene treatment (P-value for the overlap using a hypergeometric test: p < 4.657*10^-143^) (Figure S3A). Notably, more than 90% of these common DEGs retained the directionality of their expression change (i.e. genes upregulated by MBZ were also upregulated by ethylene treatment and vice versa) (Figure S3B, C). While this overlap was highly significant and supported the hypothesis that MBZ elicited ethylene responses, there was a high number of differentially expressed genes that didn’t overlap. We reasoned that this might be due to differences in developmental and growth conditions of our MBZ-treated plants and of those plants used for the published ethylene treatment dataset (Chang et al., 2013). For instance, we had used 14-day-old seedlings grown under long day conditions before we conducted the treatment and the published data had utilized 3-day-old, dark-grown seedlings for eliciting the classical ethylene triple response phenotype. To compare ethylene responses to MBZ responses under the same growth conditions and plant age, we performed another mRNA-seq experiment using roots of 14-day-old Arabidopsis grown on ½ MS media treated with 10 µM MBZ or 10 µM ACC (the ethylene precursor) for 3 hours. We then again compared the overlap of the responsive genes. Out of 826 DEGs upon the MBZ treatment, 524 were overlapping with 741 DEGs upon 3 hours of ACC treatment, accounting for 63 and 71% of DEGs upon MBZ and ACC treatment respectively (Figure 2C). 77% upregulated DEGs of the MBZ treatment overlapped with 65% upregulated DEGs of the ACC treatment (Figure 2D). These fractions were 54% and 75% for downregulated DEGs (Figure 2E). Taken together, our data strongly suggested that application of MBZ induces the ethylene pathway but in addition might also elicit non-ethylene related effects.

**Figure 2.**
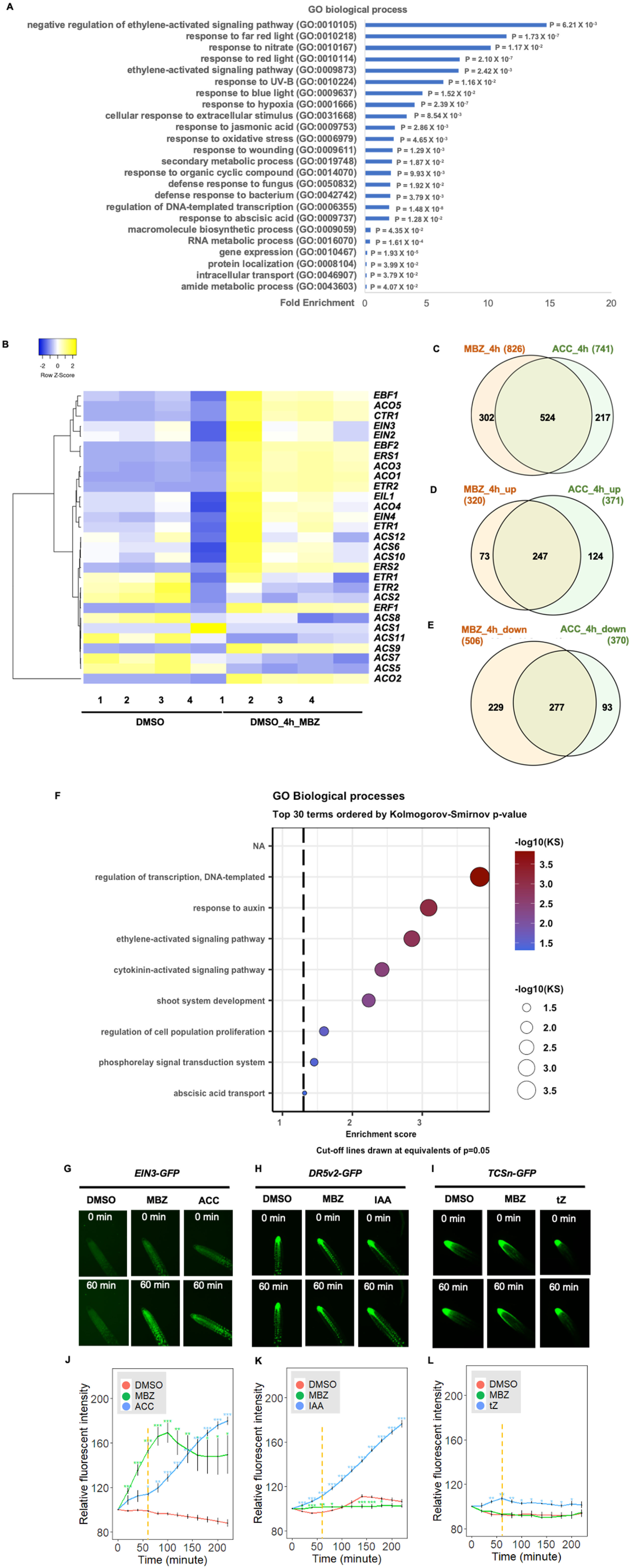
MBZ treatment regulates the ethylene pathway. (A) Gene ontology (GO) analysis upregulated genes (abs(log2) fold change > 1) from DEGs upon MBZ treatment in roots of 14-day-old Arabidopsis seedlings. (B) Heatmap of DEGs involved in the ethylene pathway, which were detected to be differentially expressed upon MBZ treatment in the RNA-seq analysis. The left 4 columns of DMSO are data from 4 replicates of samples from DMSO (Mock) plates. The right 4 columns of MBZ are data from 4 replicates of samples from MBZ treatment plates. The heatmap was made online by heatmapper (http://heatmapper.ca/expression/), The cluster method is complete linkage. Color bar indicates the relative expression level. (C-E) Venn diagrams showing differentially expressed genes in 3 h MBZ treatment and 3 h ACC treatment. All genes (FDR < 0.05; abs(log2) fold change > 1) (C), upregulated genes (FDR < 0.05; log2 fold change > 1) (D), downregulated genes (log2 fold change <-1) (E). (F) Dot plots showing the most enriched GO terms with KS p-value < 0.05 from DEGs upon MBZ treatment in lateral root tips of 14-day-old seedlings. (G-I) Representative images of lateral root tips of two-week-old plants in response to negative control, positive control and MBZ treatment using ethylene (*EIN3-GFP*)(G), auxin (*DR5v2-GFP*)(H), and cytokinin (*TCSn-GFP*)(I) reporter lines at 0 and 60 minutes of treatment. (J) Relative fluorescent intensity of *EIN3-GFP* lateral root tips treated over a course of 4 h with DMSO, MBZ, and ACC. (K) Relative fluorescent intensity of *DR5v2-GFP* lateral root tips treated over a course of 4 h with DMSO, MBZ, and IAA. (L) Relative fluorescent intensity of *TCSn-GFP* lateral root tips treated over a course of 4 h with DMSO, MBZ, and tZ. 6 lateral roots for each treatment were used. Unpaired, two-tailed Student’s t-tests were used to compare the data of MBZ or the corresponding positive controls to DMSO at each timepoint. *: p<0.05; **: p<0.01; ***: p<0.001.

### MBZ induces ethylene responses

To investigate how MBZ treatment effects the ethylene response, we compared plant responses to MBZ and the ethylene precursor ACC. The most characteristic response to ethylene treatment is the triple response of etiolated seedlings (Guzman and Ecker, 1990) (Figure 3A to C), which includes shortened and thickened roots and hypocotyls, and an exaggerated apical hook. Consistent with our hypothesis that MBZ activates ethylene responses, 3-day-old etiolated seedlings treated with 10 µM MBZ showed a prominent triple response phenotype (Figure 3A to C). Using different doses of MBZ, we found that 2.5 µM of MBZ was sufficient to induce the triple response phenotype in a dark environment (Figure S3A to F), we therefore started to use 2.5 µM of MBZ to observe triple responses. Consistent with MBZ mimicking ethylene treatment, MBZ induced triple response was abolished in the ethylene insensitive mutants *ein2-5* and *ein3 eil1* (Figure S3D to F). Ethylene treatment inhibits root elongation mainly by repressing cell elongation rather than reducing the length of meristem (De Cnodder et al., 2005; Vaseva et al., 2018). We therefore measured the length of root cells of 4-day-old etiolated seedlings treated with 2.5 µM MBZ and 10 µM ACC, and found that similar to the ACC treatment, the mature cell size was reduced by MBZ treatment (Figure 3D to F). Altogether, our data strongly suggested that MBZ elicits its effects via the ethylene pathway.

**Figure 3.**
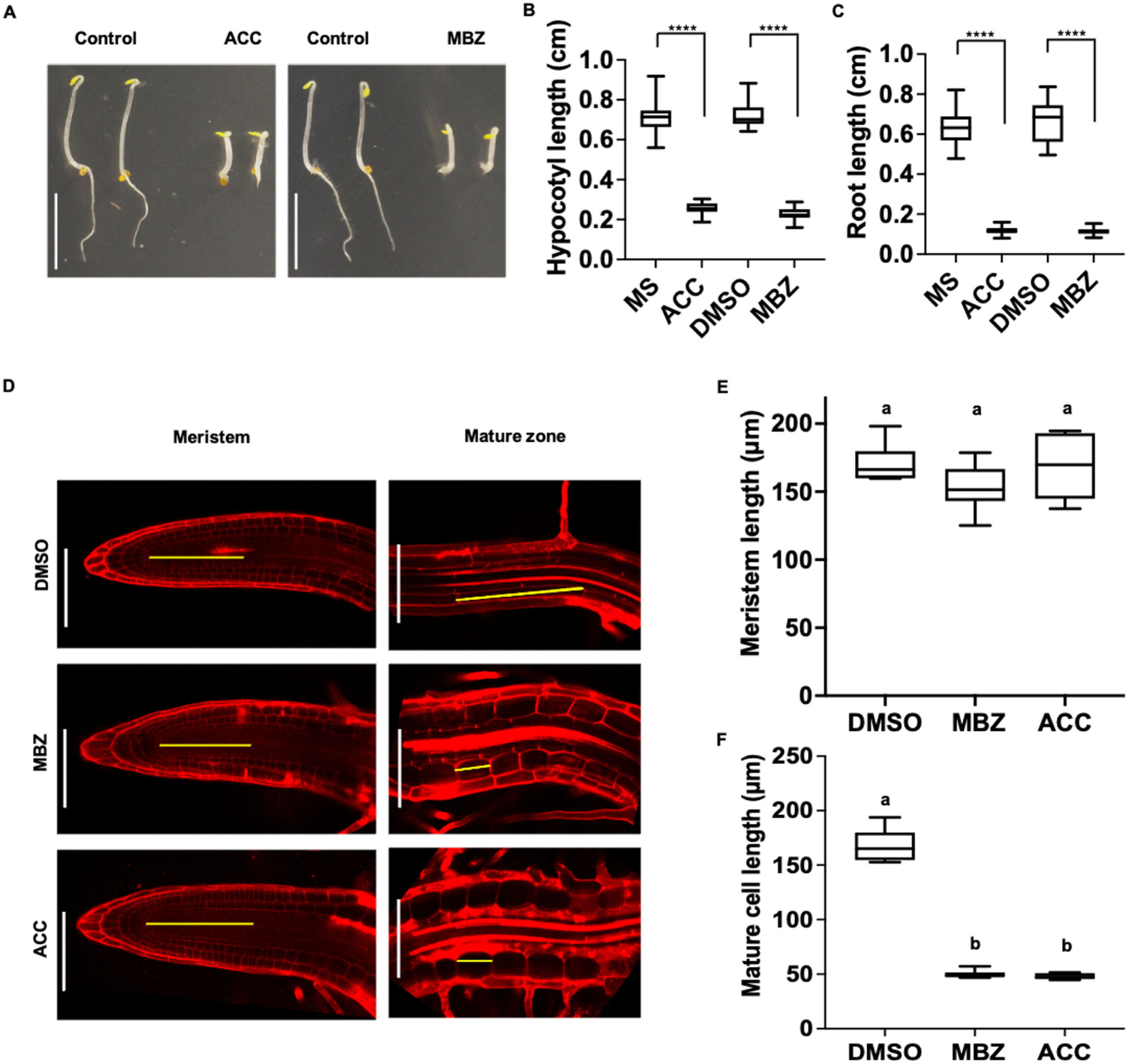
MBZ induces ethylene responses and mimics ACC treatment. (A) 4-day-old etiolated seedlings grown on MS, 10 µM ACC, DMSO, 10 µM MBZ plates. 20 Col-0 seedlings for each treatment were observed and 2 representative seedlings for each treatment are shown in the panel. (B-C) Quantification of hypocotyl length (B) and primary root length (C) of seedlings in (A). 20 Col-0 seedlings for each treatment were used. **** p < 0.0001. (D) Images of median longitudinal optical sections of primary root meristem and mature zone stained with propidium iodide (PI) of 4-day-old etiolated Col-0 on DMSO, 2.5 µM MBZ, and 10 µM ACC plates, 6-10 seedlings were observed for each treatment. (E-F) Meristem length (E) and mature cell size (F) of seedlings in (D). 6-10 seedlings for each treatment were used. Different letters label significant different values (p<0.0001). One-way ANOVA and *post hoc* Tukey testing were used for statistical analysis in (E) and (F). (Scale bar: **A**, 5 mm; **D**, 100 µm.). Boxplots: Whiskers: min/max values; hinges: 25^th^ to 75^th^ percentile; mid-line: median.

As we wanted to identify the mechanism by which MBZ changes the steepness of lateral root angles and given our data that MBZ acts on the ethylene pathway, we conducted an RNA-seq experiment on lateral root tips that had been treated either with DMSO or MBZ for 3 hours. As sample collection was very challenging and took an hour per replicate, the sampled lateral roots had been treated for a time between 3-4 hours. The most enriched gene ontology categories were “regulation of transcription”, “response to auxin”, “ethylene activated signaling pathway” and “cytokinin-signaling pathway” (Figure 2F). Given that the time window for the RNA-seq coincided with the change of lateral root angle, it was not surprising to detect auxin and cytokinin related genes, as these two hormonal pathways had been shown to be involved in determining the GSA of lateral roots (Rosquete et al., 2013a; Roychoudhry et al., 2013; Waidmann et al., 2019b). However, given the enrichment of genes assigned to these pathways we wanted to check whether these pathways are activated before or after ethylene responses were elicited by MBZ. To do this, we obtained the *DR5-GFP V2* auxin response reporter line (Liao et al., 2015a), the *35S-EIN3-GFP* ethylene reporter line in the *ein3 eil1* background (He et al., 2011), and the *TCN-GFP* cytokinin reporter line (Zurcher et al., 2013). We then conducted time-lapse spinning disk confocal microscopy on lateral root tips from 14-day-old seedlings that were either treated with a DMSO control solution, MBZ, or the respective positive control (the auxin IAA, the cytokinin Zeatin, or the ethylene precursor ACC). All positive control treatments swiftly induced the expression of the reporter genes in lateral roots (Figure 2G-L; Suppl. Movies 1, 3, 4, 6, 7, 9). Consistent with our hypothesis that MBZ induced the ethylene pathway, MBZ treatment drastically induced the ethylene reporter within the first 20 minutes after treatment, even before the ethylene precursor ACC induced the reporter (Figure 2G, J; Suppl. Movies 1-3). MBZ treatment did not lead to an increased signal of the auxin response reporter, while the DMSO control led to a slight decrease of the reporter signal after 40 minutes, and the IAA treatment led to a strong increase throughout the whole duration of the experiment (Figure 2H, K; Suppl. Movies 4-6). There was no induction of the cytokinin reporter by MBZ in the timeframe we observed the roots (Figure 2I, L; Suppl. Movies 7-9). Overall, these data strongly suggested that MBZ strongly and swiftly induces the ethylene pathway in lateral root tips.

### MBZ activates ethylene signaling

Next, we set out to locate the target of MBZ in the ethylene pathway. First, we tested whether MBZ induces ethylene biosynthesis, or acts on the perception of ethylene, or on the ethylene signaling pathway. To test whether ethylene biosynthesis is affected, we measured ethylene production by GC-MS in 3-day-old etiolated Col-0 seedlings that were treated with 10 µM MBZ or ACC. Ethylene production was dramatically induced by 10 µM ACC, but not detected in DMSO or 10 µM MBZ-treated-seedlings (Figure 4A). The results suggested that MBZ acts downstream of ethylene biosynthesis. To further test this hypothesis, we utilized gain-of-function mutants in the ethylene receptor *ETR1*. Root growth and GSA in strong *ETR1* alleles *etr1-1 and etr1-3,* were still strongly altered upon MBZ treatment (Figure 4B to E). Moreover, 4-day-old etiolated seedlings of *etr1-3* treated with 2.5 µM MBZ showed the typical triple response phenotype, demonstrating that ethylene responses are triggered by MBZ even without induced ethylene production or the perception of ethylene by ETR1 (Figure S3A to C). In contrast to the response to MBZ, *etr1-3* plants were completely insensitive to the treatment with 10 µM of ACC (Figure S3A to C). Collectively, these results show that MBZ induces ethylene responses by targeting factors downstream of ethylene biosynthesis and perception.

**Figure 4.**
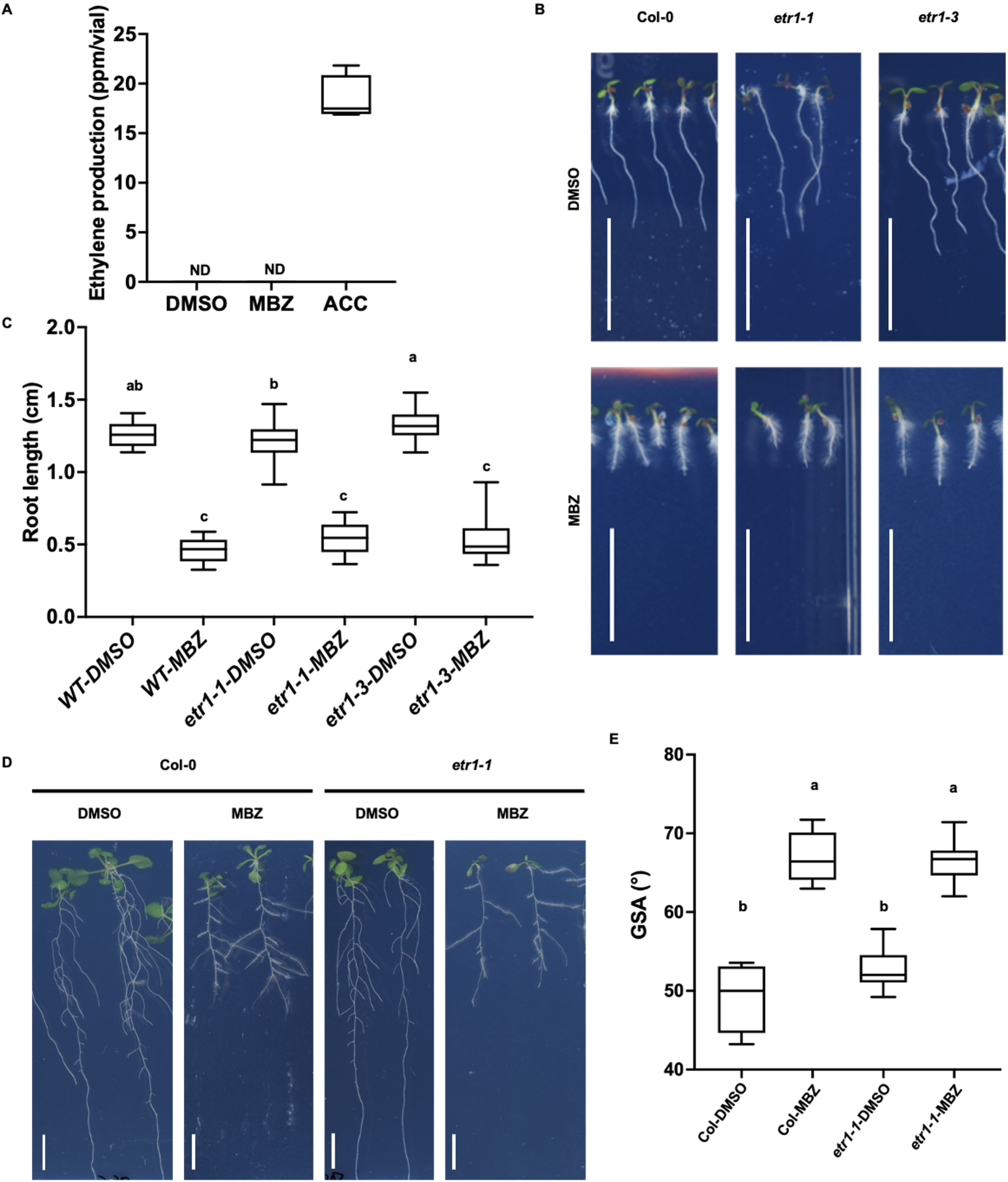
MBZ acts downstream of ethylene biosynthesis. (A) Quantification of ethylene production by GC-MS of 4-day-old etiolated seedlings of Col-0 in vials with DMSO, 10 µM MBZ, and 10 µM ACC medium. 100 seedlings for every vial, 4 biological replicates for each treatment, ND: not detectable. (B) 6-day-old seedlings of Col-0, *etr1-1*, and *etr1-3* grown on DMSO and 1.2 µM MBZ plates. 20 seedlings were observed. (C) Quantification of root length in (A). 20 seedlings were countified. (D) 16-day-old seedlings of Col-0 and *etr1-1* grown on DMSO and 1.2 µM MBZ plates. 20 seedlings were observed. (E) Quantification of GSA in (C). 20 seedlings were countified. One-way ANOVA and *post hoc* Tukey testing were used for statistical analysis in b and d. Different letters label significant different values (p<0.0001). (Scale bar: **B**, **D**, 1 cm.) Boxplots: Whiskers: min/max values; hinges: 25^th^ to 75^th^ percentile; mid-line: median.

To uncover which components of ethylene signaling are targeted by MBZ, we studied MBZ responses in key mutants of the ethylene signaling pathway. First, we checked the root inhibition phenotype and GSA in *ctr1-1* mutant treated with MBZ and compared these responses to the Col-0 wild type. *CTR1* is the first negative regulator of ethylene signaling acting downstream of ethylene receptors, and *EIN2* and *EIN3/EIL1* are positive regulators that act downstream of *CTR1* to trigger ethylene signaling (Figure 5A). Roots of Col-0 were sensitive to 1.2 µM MBZ treatment and showed short roots and increased root hair development (Figure 5B, C). However, *ctr1-1* mutants in which ethylene signaling is constitutively active showed short roots and abundant root hairs without any treatment at an early stage. Due to this pronounced early-seedling phenotype, we could not confidently assess changes in response to MBZ. However, we noticed that root growth remained insensitive to MBZ (Figure 5B, C). We further measured GSA at a later stage of development, which we had found to be a major effect of MBZ treatment. 3-week-old *ctr1-1* plants were insensitive to MBZ with regards to GSA (Figure 5D, E). Interestingly, *ctr1-1* without MBZ treatment showed an increased lateral root angle phenotype, which mimics MBZ treatment, suggesting that CTR1 might be the target of MBZ in ethylene signaling. Consistent with this hypothesis, root growth and GSA of *ein2-5* and *ein3 eil1* was clearly not affected by MBZ treatment (Figure 5B, C, F, G) and in fact, *ein2-5* mutant plants, which are insensitive to ethylene, displayed steeper lateral root angles under control conditions compared to the wildtype (Figure 5F, G). We further tested triple responses in *ctr1-1*, *ein2-5* and *ein3 eil1* mutants. Consistent with the observation of root inhibition and GSA in *ctr1-1*, these plants showed a constitutive triple response regardless of MBZ treatment (Figure S3D to F). *ein2-5* and *ein3 eil1* plants were also insensitive to MBZ treatment (Figure S3D to F). Taken together our data showed that MBZ activates ethylene signaling most likely by interfering with CTR1 in the ethylene signaling pathway.

**Figure 5.**
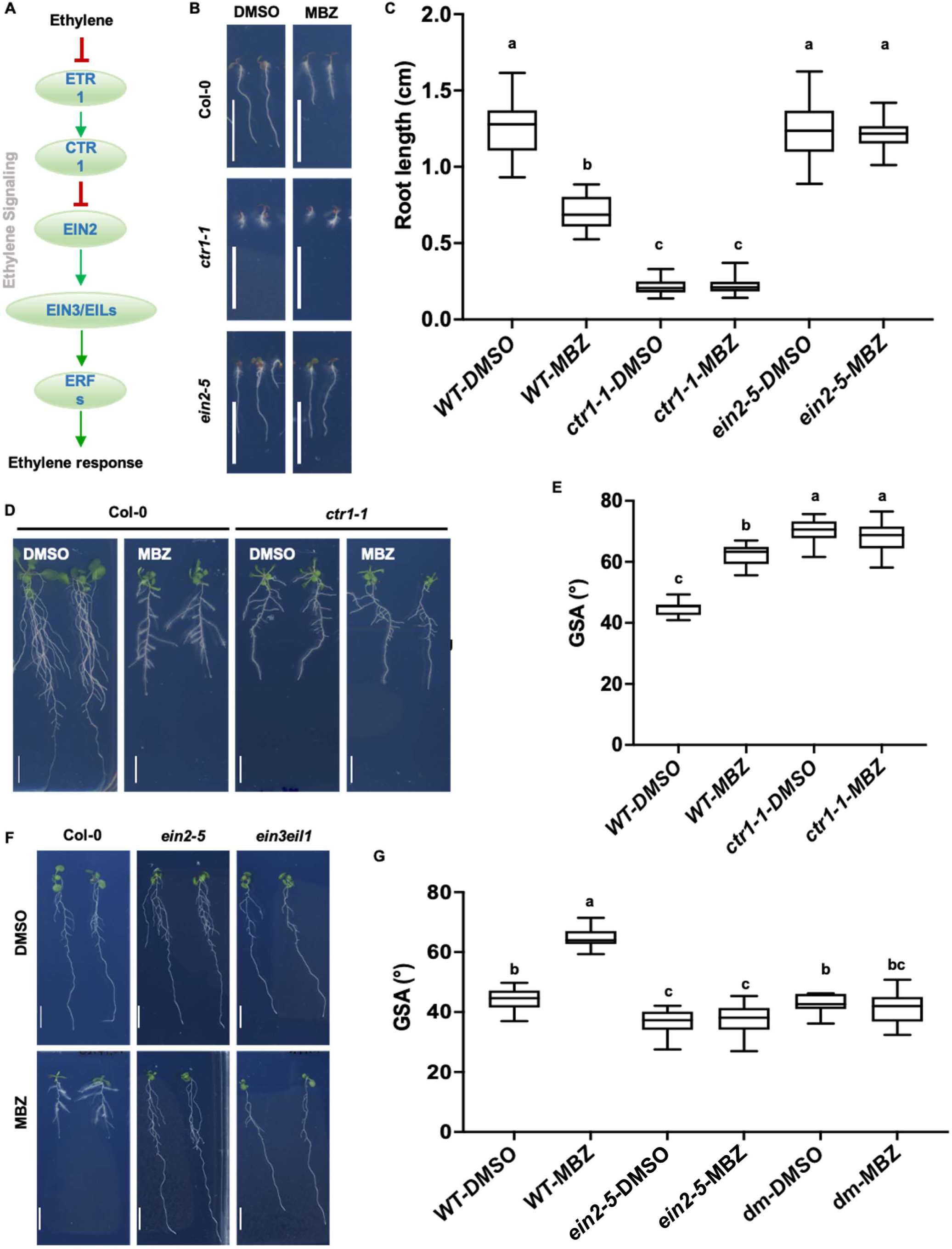
MBZ targets the ethylene signaling pathway. (A) Diagram of ethylene signaling pathway. (B) 7-day-old seedlings of Col-0, *ctr1-1*, and *ein2-5* grown on DMSO and 1.2 µM MBZ plates. (C) Quantification of primary root length of seedlings in (B) 15-20 seedlings were quantified. Similar results were obtained from three biological replicates of the experiment. (D) 19-day-old seedlings of Col-0 and *ctr1-1* grown on DMSO and 1.2 µM MBZ plates. (E) Quantification of GSA of seedlings in (D). 10 seedlings were quantified. Similar results were obtained from three biological replicates of the experiment. (F) 16-day-old seedlings of Col-0 and *ein2-5*, *ein3 eil1* grown on DMSO and 1.2 µM MBZ plates. (G) Quantification of GSA of seedlings in (F). *dm* stands for *ein3 eil1* double mutants. One-way ANOVA and *post hoc* Tukey testing were used for statistical analysis in **C**, **E**, and **G**. Different letters label significant different values (p<0.0001). (Scale bar: **B**, **D**, **F**, 1 cm.). Boxplots: Whiskers: min/max values; hinges: 25^th^ to 75^th^ percentile; mid-line: median.

### MBZ is a potent inhibitor of CTR1 kinase activity

To pinpoint whether CTR1 is the target of MBZ, we capitalized on what is known about the activity of MBZ in mammalian systems. There, MBZ has been reported to be an inhibitor of the serine/threonine kinase *homo sapiens* (*hs*) MAPK14 (Ariey-Bonnet et al., 2020). Because CTR1 is also a serine/threonine protein kinase (Kieber et al., 1993), we hypothesized that CTR1 is the direct target of MBZ in plants. Since it was shown that there are four critical amino acid residues in the catalytic site of *hs*MAPK14 that are relevant for MBZ binding (Ariey-Bonnet et al., 2020), we first tested *in silico* whether these residues were conserved between *hs*MAPK14 and CTR1. For this, we aligned the amino acid sequence of the Arabidopsis CTR1 and *hs*MAPK14 kinase domains (Figure 6A, Figure S4A, B). We found three (Lys 578, Thr 625, and Asp 694) of the four critical amino acid residues were conserved between the CTR1 kinase domain and that of *hs*MAPK14 (Figure 6A). The 4th critical residue is Met 109 in *hs*MAPK14, which corresponds to Leu 628 in CTR1. Because both Met and Leu are non-polar amino acids with hydrophobic side chains and similar structure, it appeared possible that the replacement of Met by Leu in the CTR1 kinase domain does not significantly change the pocket structure for MBZ binding. We therefore hypothesized that the kinase domains of CTR1 and MAPK14 might have similar structures and function with regards to binding MBZ (Figure S4B).

**Figure 6.**
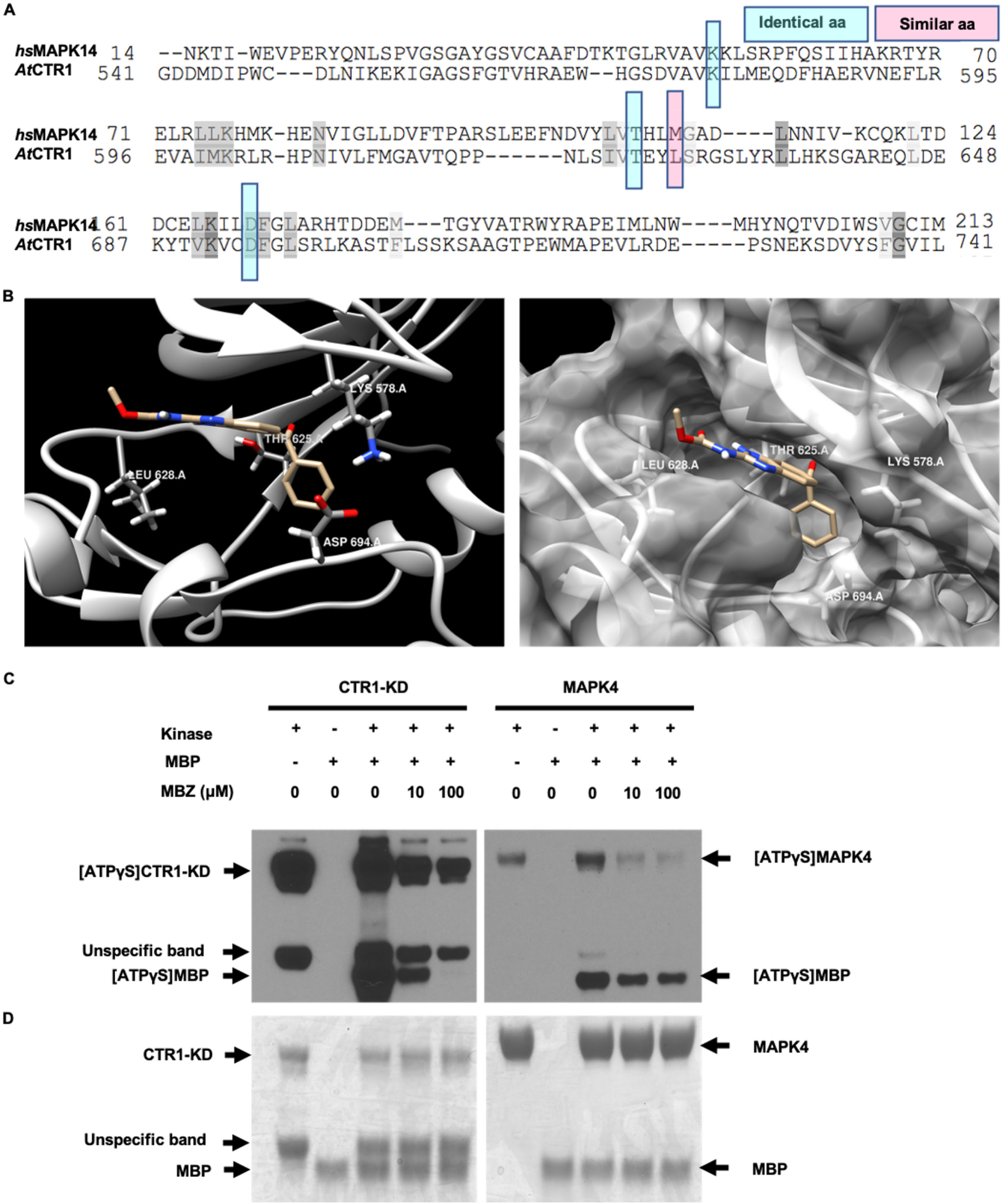
MBZ inhibits CTR1 kinase activity. (A) Sequence alignment of *hs*MAPK14 and CTR1-KD in Arabidopsis around 4 core amino acids (Highlighted by blue and pink frames) for MBZ binding. (B) Molecular modeling of the interaction between MBZ (coppery) and CTR1-KD (silvery). In the left panel, the key residues that contribute to the binding with CTR1-KD were highlighted. The right panel showed the surface (gray) of the catalytic pocket of CTR1- KD. (C) Western blot using Recombinant Anti-Thiophosphate ester antibody under different concentration of MBZ treatment. Left panel: CTR1-KD; Right panel: MAPK4. MBP is the substrate in both panels, MBZ concentration is 0, 10, and 100 µM as labeled in different reactions, [ATPγS]CTR1-KD: ATPγS binding CTR1-KD, [ATPγS]MAPK4: ATPγS binding MAPK4, MBP that obtained ATPγS from CTR1-KD or MAPK4, which is labeled as [ATPγS]MBP. Note: ATPγS: Adenosine 5’-(3-thiotriphosphate) tetralithium salt (Sigma, Cat# A1388) (D) Coomassie blue staining of the gel shown in (C). CTR1: ∼100 ng, MAPK4: ∼1 µg. Myelin Basic Protein (MBP, a common kinase substrate, Sigma Cat. M1891): 2 µg, ATPγS: 1 mM. MBZ: 0, 10, or 100 µM. Arrows label the position of these bands on the gel.

To test this hypothesis, we first performed an *in silico* docking analysis using the published crystal structures of CTR1 (Mayerhofer et al., 2012)(PDB: 3PPZ) and MBZ (CHEMBL685). We found that MBZ is predicted to bind into the pocket of the CTR1 kinase domain (Figure 6B) formed by the aforementioned key amino acids (Figure 6A, B, Figure S4B). Overall, these analyses indicated that CTR1 could be the target of MBZ in Arabidopsis.

To further test this model, we expressed the recombinant kinase domains (KD) of CTR1 and MAPK4 (a MAPK protein in Arabidopsis with a highly similar amino acid sequence to *hs*MAPK14) (Table S1, Figure S4C, D). We then confirmed the kinase inhibitory activity of MBZ through an *in vitro* phosphorylation assay via anti-Thiophosphate-Ester antibodies (Allen et al., 2007). Using the phosphorylation state of MBP (substrate) as the readout of the kinase activity, we found that MBZ inhibited the kinase activities of both CTR1-KD and MPK4 in a concentration-dependent manner, however, the inhibition for CTR1-KD was much more pronounced (Figure 6C, D). In particular, 10 µM MBZ strongly inhibited the kinase activity of CTR1-KD, while 100 µM MBZ totally blocked its activity. This assay demonstrated that MBZ can directly inhibit the kinase activity of CTR1-KD. Together with our genetic data that *ctr1-1* mutant mimicked the effect of MBZ on GSA and the triple response phenotypes (Figure 5D, E, Figure S3D-F), and our molecular docking data (Figure 6B), our data strongly suggest that CTR1 is a direct target of MBZ in Arabidopsis. To further test the hypothesis that MBZ acts upon CTR1, we utilized a readout directly downstream of CTR1. Previous studies had shown that functional CTR1 phosphorylates the carboxyl end of EIN2, leading to its degradation (Ju et al., 2012; Qiao et al., 2012a; Wen et al., 2012). When ethylene inactivates CTR1, EIN2-CEND is stabilized and moves into the nucleus to activate downstream ethylene responses. We therefore tested if MBZ treatment mimicked ethylene for inducing EIN2 accumulation in the nucleus using 3-day-old etiolated seedlings of two independent transgenic *EIN2-GFP* reporter lines in which the GFP was fused to EIN2’s C-terminus (*pEIN2::EIN2-GFP* in the *ein2-5* background (Fu et al., 2021)). As expected, treatment with ACC quickly increased the nuclear accumulation of EIN2-GFP in root cells (Figure 7A, B). Consistent with our hypothesis that MBZ acted on CTR1, MBZ also significantly increased accumulation of EIN2-GFP in the nucleus (Figure 7A, B). Taken together, our data support a model in which MBZ is a small molecular inhibitor of the CTR1 kinase activity and thereby triggers the ethylene signaling pathway by permitting C-terminal of EIN2 translocation into the nucleus, which in turn triggers ethylene downstream responses (Figure 7C, D).

**Figure 7.**
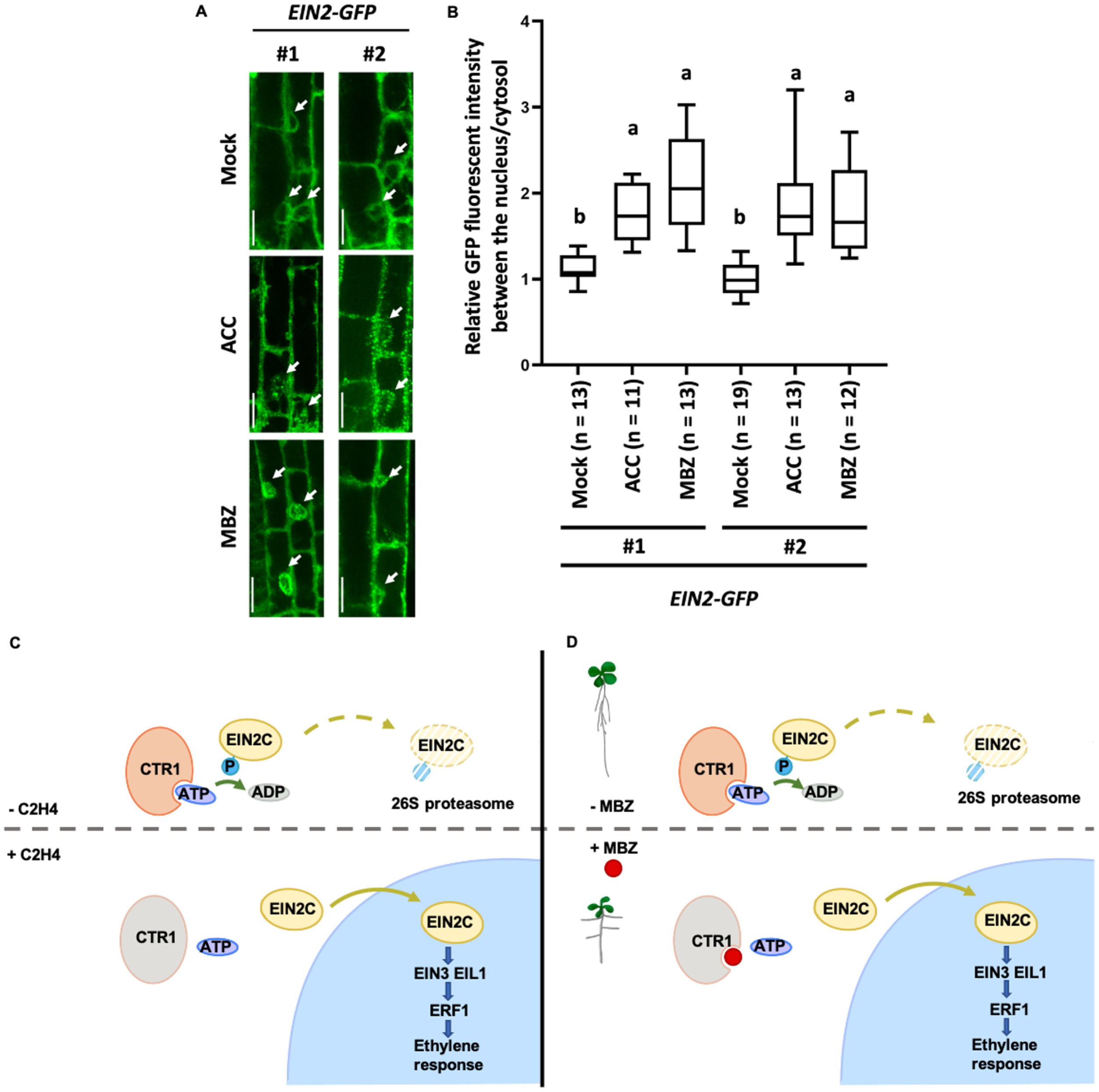
MBZ activates ethylene signaling. (A) Different patterns of nuclear accumulation of *EIN2-GFP* in root cells. 3-day-old etiolated seedlings treated with Mock, ACC (100 µM), or MBZ (10 µM) for 3 hours. Arrows indicate nuclei. (Scale bar: 20 µm). (B) Quantification of Relative GFP fluorescent intensity between the nucleus and cytosol of *EIN2-GFP* in **A**. Boxplots: Whiskers: min/max values; hinges: 25^th^ to 75^th^ percentile; mid-line: median. One-way ANOVA and *post hoc* Tukey testing were used for statistical analysis. (C,D) Model for MBZ activation of ethylene signaling. (C) Ethylene suppresses CTR1 activity to promote EIN2C translocation and activates ethylene signaling. (D) MBZ inhibits CTR1 activity by binding to its kinase domain and promotes EIN2C translocation to activate the ethylene signaling. Note: EIN2C: C-terminal of EIN2; ATP: Adenosine triphosphate; ADP: adenosine diphosphate; C2H4: ethylene; P: Phosphate group. The CTR1 with orange color means active CTR1 proteins and the CTR1 with gray color means inactive CTR1.

## DISCUSSION

Our chemical genetic screen led to the identification of MBZ as a potent inhibitor of the kinase activity of CTR1, which is a negative modulator of the ethylene signaling pathway (Figure 7C). MBZ therefore is an activator of ethylene signaling, and to our knowledge the first small molecule that acts downstream of ethylene receptors and that directly activates ethylene signaling. In previous studies, ethylene gas and ACC have been used to activate the ethylene signaling pathway. However, there were some disadvantages to use these approaches. In the case of treatment with ethylene gas, it is difficult to maintain a continuous treatment over longer periods of time due to the technical challenges for constantly maintaining the required ethylene concentrations. Because its controlled application is technically less challenging, ACC has been used more widely to study the role of ethylene. However, ACC was recently found to be a signaling molecule which can act independently of the ethylene pathway (Mou et al., 2020). Moreover, the necessity for enzymatic ACC conversion to ethylene in plants might be limiting in different tissues or conditions. While MBZ is not a perfect ethylene substitute and might have unspecific effects beyond activating ethylene signaling, as a direct activator of the ethylene signaling pathway, it promises to be a useful tool for the field. Such tools have been lacking so far despite several attempts that have been made to identify small molecular regulators of ethylene signaling. Neither approaches targeting ethylene biogenesis and perception, nor approaches targeting downstream auxin biosynthesis had yielded in molecules acting on ethylene signaling (He et al., 2011; Li et al., 2017; Sun et al., 2017; Zhu et al., 2019).

Because CTR1 contains the critical MBZ binding residues that had been elucidated in the human MBZ target *hs*MPK14, and because of the profound genetic evidence for CTR1, it is somewhat unlikely that other components in the ethylene pathway such as the E3 ligases ETP1/2 and EBF1/2, which negatively regulate EIN2 and EIN3 respectively (Guo and Ecker, 2003; Qiao et al., 2012b), are targets of MBZ. Another signaling pathway that impinges on ethylene signaling is the glucose-activated TOR signaling pathway, which had recently been shown to regulate EIN2 activity through phosphorylation (Fu et al., 2021). However, the TOR kinase does not contain any of the 4 key residues known for MBZ binding. Moreover, unlike MBZ, none of the inhibitors of TOR, including Rap, Torin2, and AMA, did induce the ethylene induced triple response phenotype (Fu et al., 2021). In addition, *ein2-5* and *ein3 eil1* mutants displayed a shorter hypocotyl when the Glu-TOR- EIN2 pathway was inhibited (Fu et al Nature 2021, Figure 1b), but these mutants did not respond to MBZ treatment (Figure S3D, E). Altogether, it is most likely that the observed response of MBZ on the ethylene signaling is exerted via its binding to CTR1.

While most of the effects of MBZ are related to its activation of ethylene signaling, our RNA profiling indicated additional effects of MBZ. Consistently, treatment with high concentration of MBZ or long-term treatment with it led to strongly inhibited plant growth and development, including a disrupted root apical meristem and morphological changes of leaves. These phenotypes have not been observed upon ethylene treatment (Figure S5A, B), raising the possibility that MBZ may have other targets in plants. MAPK4 has the highest amino acid sequence similarity to the human MBZ target *hs*MAPK14 (Table S1).

Compared to CTR1, the effect of MBZ on MAPK4 kinase activity was much less pronounced (Figure 6C), consistent with the lack of conservation of key amino acids relevant for MBZ binding (Figure S4C). Although MBZ couldn’t inhibit the kinase activity of MAPK4 as strongly as that of CTR1-KD, a decrease of the weak MAPK4 autophosphorylation band might indicate that MBZ treatment can disrupt MAPK4 autophosphorylation (Figure 6C). This raises the possibility that MBZ might alter autophosphorylation of MAPK4 and probably other CTR1-like kinases, based on the sequence conservation of the four critical residues (Figure 6A, Figure S4B). This might explain the additional effects of MBZ. Additionally, there are a number of CTR1 paralogs in Arabidopsis. This includes the kinases encoded by AT4G24480, AT1G08720, At1g18160, AT1G67920, and AT5G11850. These kinases contain all four critical amino acids, raising the possibility that MBZ may also bind to these paralogs. Another explanation for the observed effects of MBZ that go beyond observed ethylene effects, might be that continuous activation of ethylene signaling via CTR1 might have more complicated or compounded effects than are elicited by external ethylene or ACC treatment (Figure S5C). Support for this comes from the *ctr1-1* mutant, which is dwarfed and shows aberration in leaf shape and thereby displays much stronger phenotypes than elicited with ACC or ethylene treatment (Kieber et al., 1993) (Figure 5B, D, Figure S5C). While MBZ’s predominant effect on ethylene signaling via CTR1 is strongly supported by our study, further studies will be needed to elucidate the details of MBZ specificity.

One of the most intriguing effects of MBZ on the plant is its potent ability to change GSA. This suggests that ethylene signaling is involved in the regulation of the gravitropic setpoint angle in lateral roots. While important regulatory roles for GSA are known for auxin and cytokinin, and there can be complex interplays between auxin, cytokinin and ethylene, the induction of the ethylene signaling pathway clearly dominated responses in lateral roots after they were treated with MBZ (Figure 2). This activation of the ethylene responses occurred before angles of lateral roots were changing and was maintained throughout the timescale at which GSA was changed by the MBZ treatment, which occurred after 1-4 hours of MBZ treatment (Figure 1E, F, Figure 2, Figure S2I). In our study, we found that MBZ didn’t induce ethylene biosynthesis (Figure 4A). ACC treatment had a weak but distinct effect on root angle and displayed concentration-dependent effects when applying a large range of concentrations (Figure S5C). Strong direct evidence for the impact of ethylene signaling in regulating lateral root set point angle is provided by the RSA phenotype of the *ctr1-1* mutant. In *ctr1-1*, ethylene signaling is constitutively activated and leads to an increased lateral root angle (Figure 5D, E). Conversely, we observed that the ethylene insensitive *ein2-5* mutant displays steeper lateral roots (Figure 5F, G). Overall, this exposes a yet unknown role of ethylene signaling in the regulation of lateral root set point angle. One reason why the role of ethylene in the regulation of lateral root angles hasn’t been recognized so far might be that it reduces lateral root formation by itself (Negi et al., 2008) making the study of lateral root angles more difficult. Our study strongly suggests that it will be worthwhile to explore the role of ethylene and its interaction with auxin and cytokinin in root gravitropism and RSA. Moreover, MBZ might also be used as a versatile tool for exploring the molecular mechanisms of root gravitropism and lateral root set point angles since as a small molecule, and due to its reversible mode of action, it can be easily applied to elicit controlled changes of lateral root angles. It might also be useful to reveal the mechanisms underlying GSA regulation by phytohormones and known GSA signals, such as the *LAZY* family genes (Furutani and Morita, 2021). Finally, since ethylene signaling is a widely conserved process in land plants, exploring the ability of MBZ or genetic perturbation of the ethylene pathway to change RSA in crop species might be highly worthwhile for RSA engineering.

## Acknowledgements

The research was supported by start-up funds from the Salk Institute for Biological Studies and funds from the Harnessing Plants Initiative to WB, a Salk Women & Science Special Award and a Pioneer Fund Postdoctoral Scholar Award to WH. We are grateful to Dr. Glenn R. Hicks and Dr. Natasha Raikhel for sharing the SP2000 small molecule library. We thank Dr. Cris Argueso, Dr. Dolf Weijers, and Dr. Liwen Fu for sharing seeds. We also thank Dr. Joseph R. Ecker and Dr. Joanne Chory for critical discussions. We thank the Next Generation Sequencing Core of Salk institute for NGS Library preparation & sequencing. We also thank Matthieu Platre, Nicole Gibbs, Anna Malolepszy, Björn Willige and Travis Lee for fruitful discussions and suggestions.

## Author contributions

Wenrong He and Wolfgang Busch conceived the project and the experiments. Wenrong He performed the SP2000 library screen and analysis with help from Yingnan Hou. Wenrong He performed the phenotyping, qPCR, alignment, docking, protein purification, kinase activity assay. Wenrong He conducted the microscopy and mRNA-seq on primary roots with help from Kaizhen Zhong. Hai An Truong performed the time lapse reporter analysis using spinning disk microscopy, as well as the whole root system and lateral root RNA-seq experiments. Ling Zhang conducted the mRNA-seq analysis and the statistical analysis. Wenrong He conducted heatmap analysis of mRNA-seq. Wenrong He and Ling Zhang conducted Gene ontology analysis. Min Cao provided the protocol and suggestions for kinase activity assay. Neal Arakawa performed the GC test. Yao Xiao performed the FPLC. Wenrong He and Wolfgang Busch wrote the manuscript with input from all authors. Wolfgang Busch provided materials, funding, supervision, guidance and advice on the project, the experiments, and the analysis of the results.

---

**Figure S1.**
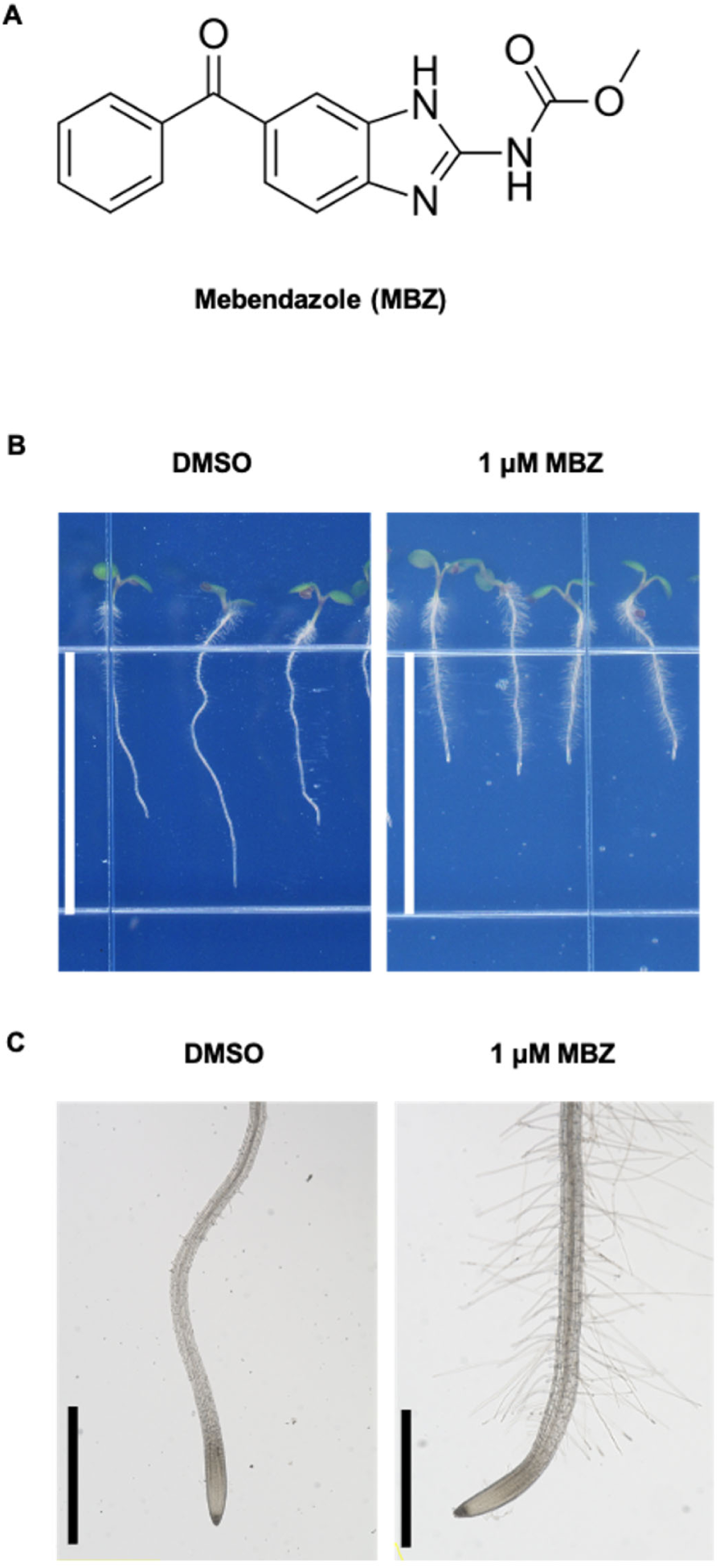
MBZ treatment affects root growth and development. (A) Chemical structure of MBZ. (B-C) Root hair phenotype of 7-day-old seedlings on DMSO and 1 µM MBZ plates recorded by a scanner (B) or a microscope (C). 20 seedlings were observed on left panel and 10 seedlings were observed on right panel. (Scale bar: **B**, 13 mm; **C**, 1000µm)

**Figure S2.**
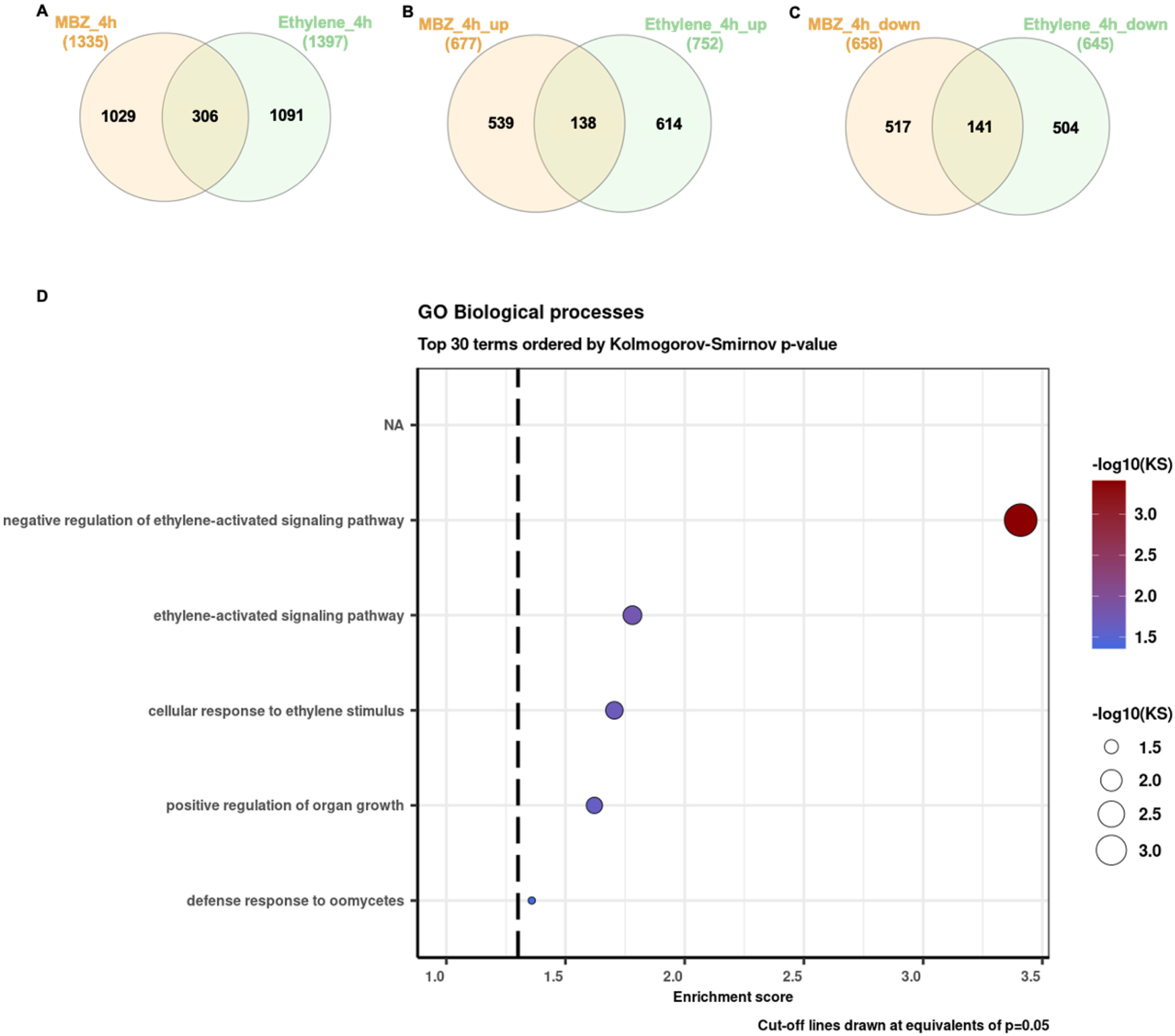
Analysis of differentially expressed genes (DEGs) upon MBZ treatment. (A-C) Venn diagrams showing differentially expressed genes in 4 h MBZ treatment in roots of 14-day-old green seedlings and published 4 h ethylene treatment in 3-day-old etiolated seedlings. All genes (FDR<0.05; abs(log2) fold change > 1) (A), upregulated genes (FDR<0.05; log2 fold change > 1) (B), downregulated genes (FDR<0.05; log2 fold change < -1) (C). (D) Gene ontology (GO) analysis of upregulated genes (FDR<0.05; log2 fold change > 1) from DEGs upon MBZ treatment.

**Figure S3.**
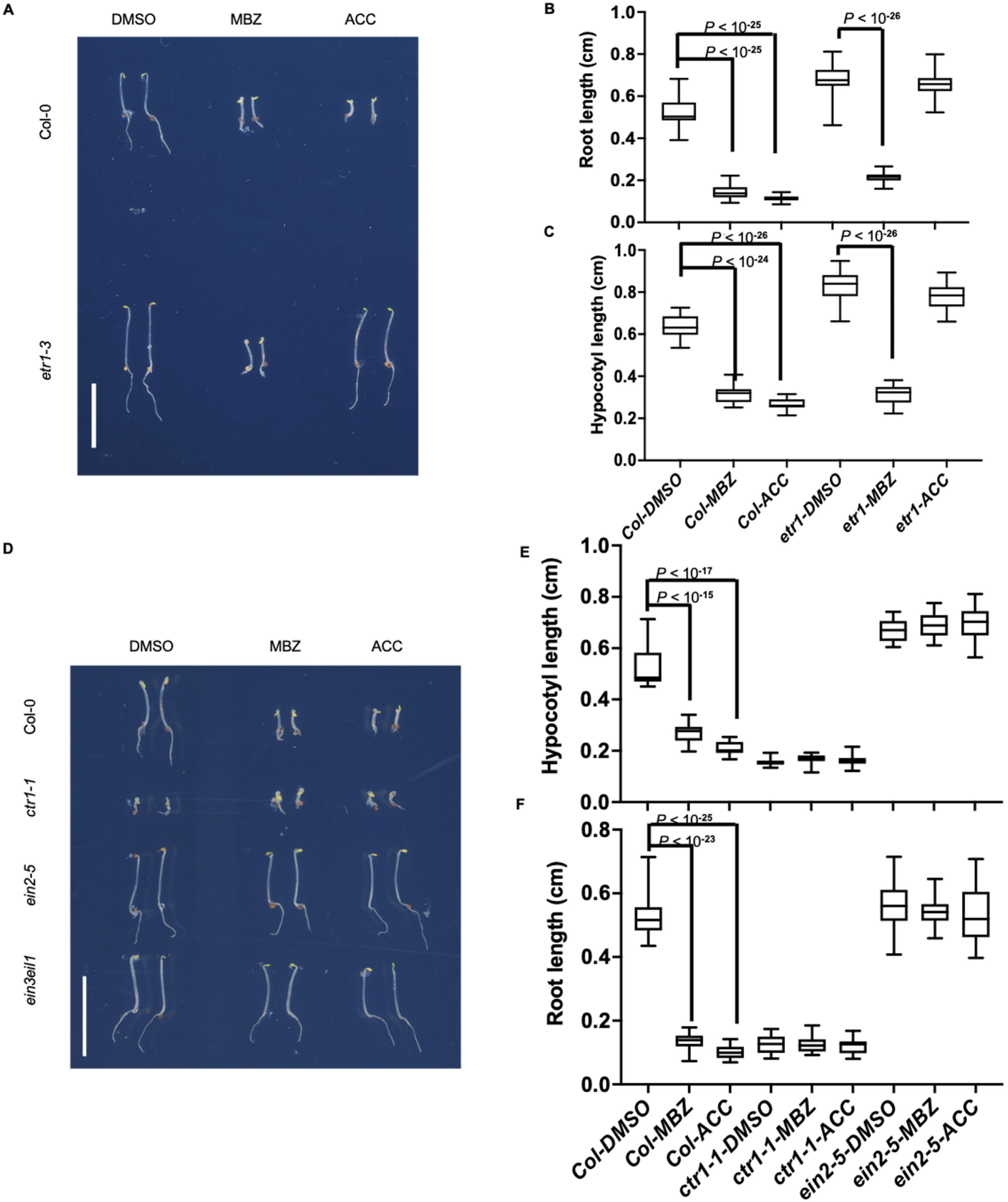
Etiolated seedling phenotypes in mutants of the ethylene pathway. (A) 4-day-old etiolated seedlings of Col-0 and *etr1-3* grown on DMSO, 2.5 µM MBZ, and 10 µM ACC plates. (B-C) Quantification of hypocotyl length (B) and root length (C) of seedlings in (A). 20 seedlings were quantified. (D) 4-day-old etiolated seedlings of Col-0, *ctr1-1, ein2-5*, and *ein3 eil1* grown on DMSO, 2.5 µM MBZ, and 10 µM ACC plates. (E-F) Quantification of hypocotyl length (E) and root length (F) of seedlings in **D**. 20 seedlings were quantified. Statistical analysis in **B, C** and **E, F** were done using unpaired, two-tailed Student’s t-tests. P-values are indicated in the figure. (Scale bar: **A, D**, 1 cm). Whiskers: min/max values; hinges: 25^th^ to 75^th^ percentile; mid-line: median.

**Figure S4.**
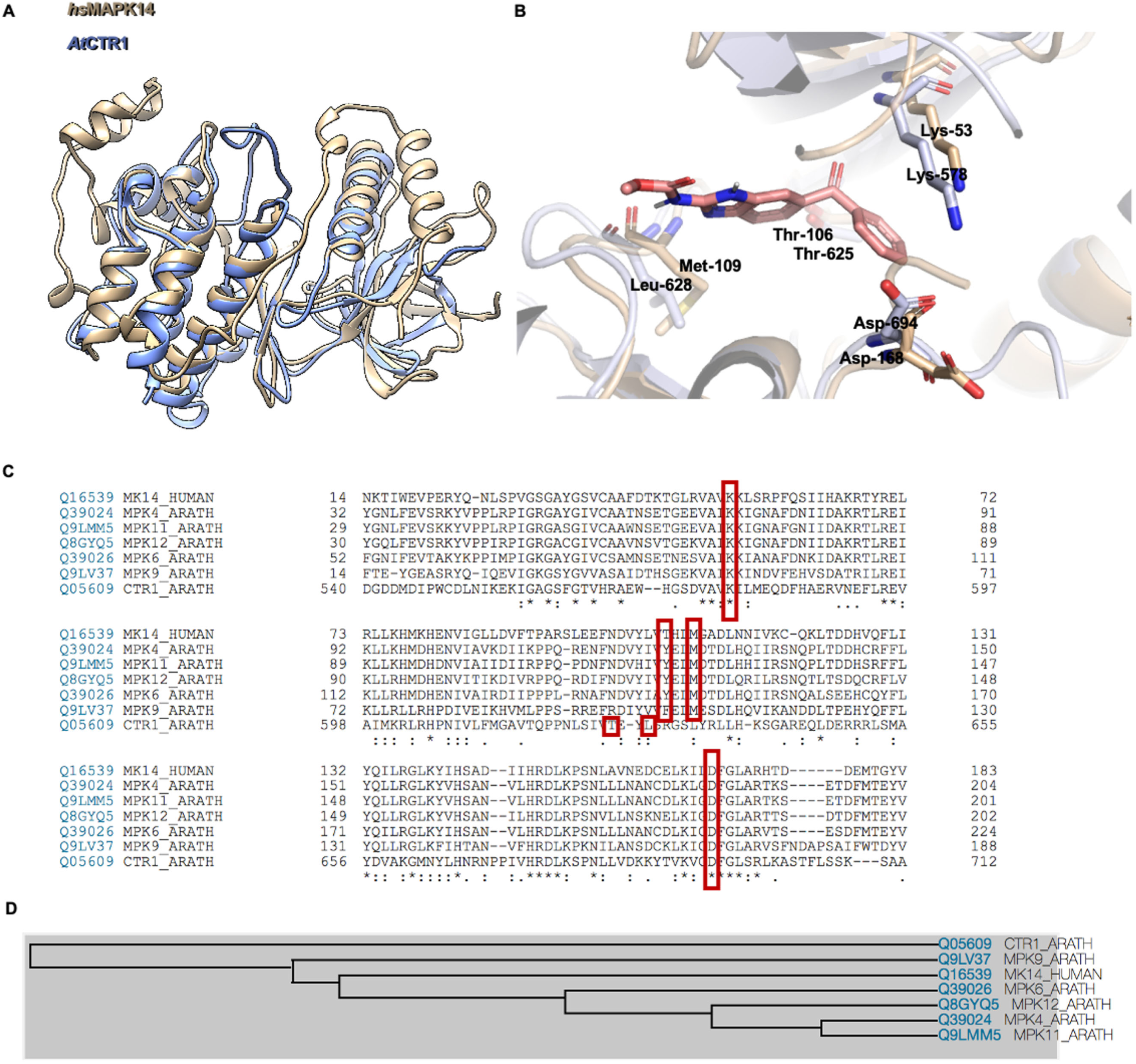
Alignment of CTR1-KD in Arabidopsis and MAPK14 in human. (A) The structure alignment of *hs*MAPK14 (coppery) and CTR1-KD (celeste). (B) The protein binding sites in *hs*MAPK14 (coppery) and CTR1-KD (periwinkle) with highlighted key residues. (C) Sequence alignment of kinase domains. Key residues in MAPK14_human and CTR1- KD in Arabidopsis are indicated by red boxes. (D) Phylogenetic analysis of kinases aligned in (C).

**Figure S5.**
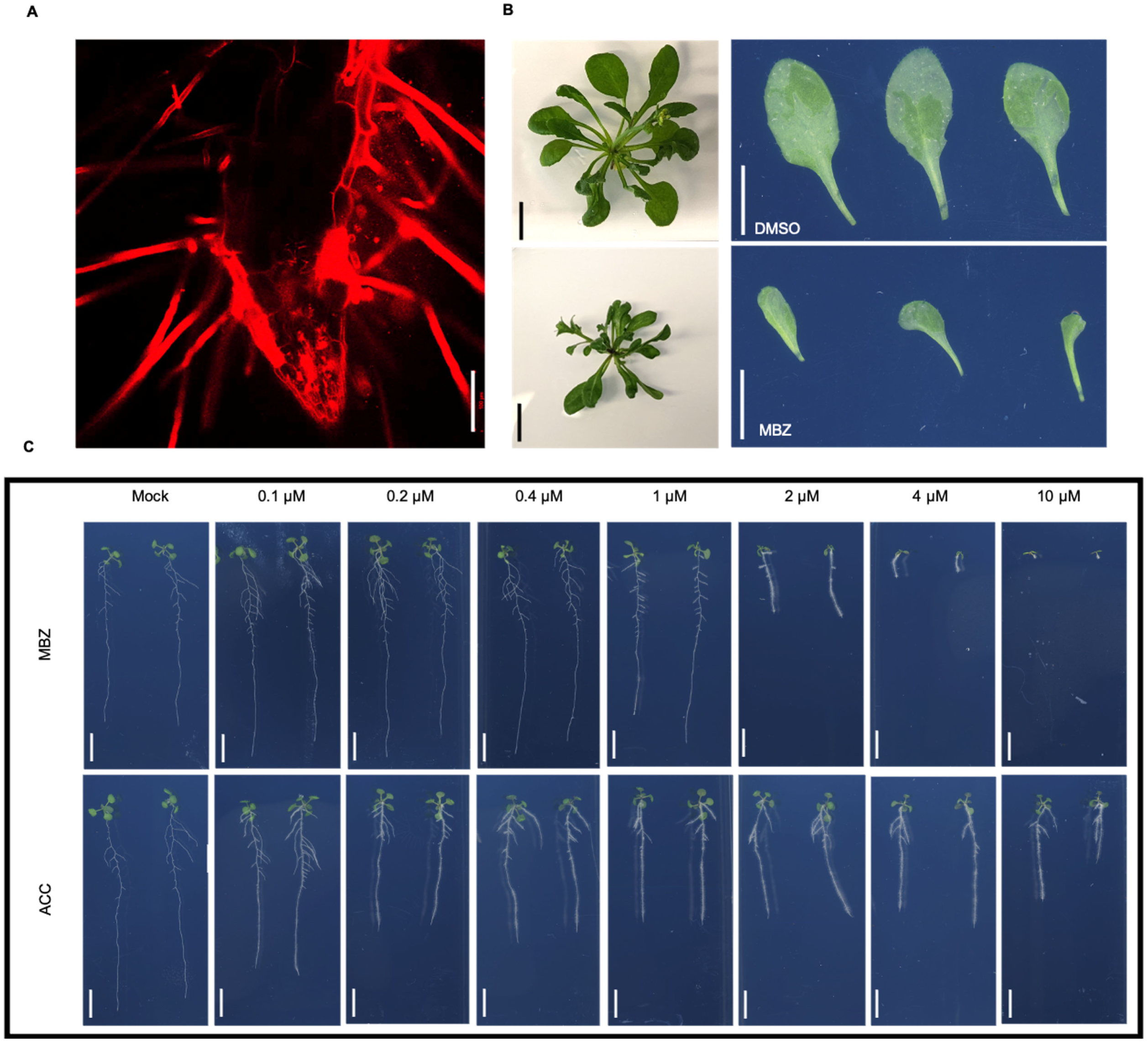
High concentrations or long-term treatment of MBZ induce phenotypes that go beyond ethylene induced effects. (A) Confocal microscopy images of a root meristem of 4-day-old etiolated WT seedling grown on 10 µM MBZ plate. (B) 38-day-old Col-0 plants grown on DMSO and 1.2 µM MBZ plates. (C) 14-day-old seedlings of Col-0 grown on increasing concentrations MBZ and ACC (0.1, 0.2, 0.4, 1, 2, 4, 10 µM) 20 seedlings were observed. Similar results were obtained from three biological replicates of the experiment. (Scale bar: **A**, 100 µm, **B**, **C**, 1 cm)

**Table S1.** The list of blast hits in Arabidopsis using the MAPK14_human sequence as BLASTP.

JVD9H54701R-Alignment-Descriptions
| Description | Scientific Name | Max Score | Total Score | Query Cover | E value | Per. ident | Acc. Len | Accession |
| --- | --- | --- | --- | --- | --- | --- | --- | --- |
| MAP kinase 4 [Arabidopsis thaliana] | Arabidopsis thaliana | 346 | 346 | 93% | 3E-117 | 51.18 | 376 | NP_192046.1 |
| mitogen-activated protein kinase 4 [Arabidopsis thaliana] | Arabidopsis thaliana | 346 | 346 | 93% | 3E-117 | 51.18 | 376 | ABR46146.1 |
| mitogen-activated protein kinase 4 [Arabidopsis thaliana] | Arabidopsis thaliana | 345 | 345 | 93% | 8E-117 | 50.88 | 376 | ABR46148.1 |
| MAP kinase [Arabidopsis thaliana] | Arabidopsis thaliana | 345 | 345 | 93% | 9E-117 | 50.88 | 376 | BAA04867.1 |
| mitogen-activated protein kinase 4 [Arabidopsis thaliana] | Arabidopsis thaliana | 344 | 344 | 93% | 1E-116 | 50.88 | 376 | ABR46162.1 |
| unnamed protein product [Arabidopsis thaliana] | Arabidopsis thaliana | 340 | 340 | 95% | 8E-115 | 47.71 | 376 | VYS62299.1 |
| MPK5 [Arabidopsis thaliana] | Arabidopsis thaliana | 339 | 339 | 95% | 9E-115 | 47.71 | 376 | OAO98154.1 |
| MAP kinase 11 [Arabidopsis thaliana] | Arabidopsis thaliana | 337 | 337 | 93% | 6E-114 | 49.71 | 369 | NP_001117210.1 |
| unnamed protein product [Arabidopsis thaliana] | Arabidopsis thaliana | 337 | 337 | 93% | 6E-114 | 49.71 | 369 | CAA0154293.1 |
| MAP kinase 5 [Arabidopsis thaliana] | Arabidopsis thaliana | 337 | 337 | 95% | 8E-114 | 47.43 | 376 | NP_567378.4 |
| unnamed protein product [Arabidopsis thaliana] | Arabidopsis thaliana | 336 | 336 | 95% | 2E-113 | 46.84 | 370 | CAA5324945.1 |
| MAP kinase [Arabidopsis thaliana] | Arabidopsis thaliana | 335 | 335 | 95% | 2E-113 | 46.55 | 370 | BAA04866.1 |
| mitogen-activated protein kinase 3 [Arabidopsis thaliana] | Arabidopsis thaliana | 335 | 335 | 95% | 2E-113 | 46.55 | 370 | NP_190150.1 |
| MAP kinase [Arabidopsis thaliana] | Arabidopsis thaliana | 330 | 330 | 95% | 3E-111 | 46.57 | 376 | BAA04868.1 |
| mitogen-activated protein kinase 12 [Arabidopsis thaliana] | Arabidopsis thaliana | 329 | 329 | 92% | 9E-111 | 49.26 | 372 | NP_182131.2 |
| mitogen-activated protein kinase 12 [Arabidopsis thaliana] | Arabidopsis thaliana | 330 | 330 | 92% | 2E-110 | 49.26 | 406 | NP_001325006.1 |
| Mitogen-activated protein kinase 6 [Arabidopsis thaliana] | Arabidopsis thaliana | 323 | 323 | 95% | 2E-108 | 46.84 | 364 | 6DTL_A |
| Crystal structure of Arabidopsis thaliana MPK6 [Arabidopsis thaliana] | Arabidopsis thaliana | 323 | 323 | 95% | 3E-108 | 46.84 | 370 | 5CI6_A |
| unnamed protein product [Arabidopsis thaliana] | Arabidopsis thaliana | 323 | 323 | 92% | 3E-108 | 46.31 | 393 | VYS60924.1 |
| MAP kinase 6 [Arabidopsis thaliana] | Arabidopsis thaliana | 323 | 323 | 95% | 4E-108 | 46.84 | 395 | NP_181907.1 |
| MAP kinase 10 [Arabidopsis thaliana] | Arabidopsis thaliana | 323 | 323 | 92% | 4E-108 | 46.31 | 393 | NP_191538.1 |
| unnamed protein product [Arabidopsis thaliana] | Arabidopsis thaliana | 323 | 323 | 92% | 5E-108 | 46.31 | 393 | CAA0387508.1 |
| MPK10 [Arabidopsis thaliana] | Arabidopsis thaliana | 323 | 323 | 92% | 6E-108 | 46.31 | 393 | OAP05435.1 |
| unnamed protein product [Arabidopsis thaliana] | Arabidopsis thaliana | 321 | 321 | 92% | 2E-107 | 46.02 | 393 | CAD5326291.1 |
| MAP kinase [Arabidopsis thaliana] | Arabidopsis thaliana | 318 | 318 | 95% | 2E-106 | 46.29 | 373 | CAB51417.1 |
| Protein kinase superfamily protein [Arabidopsis thaliana] | Arabidopsis thaliana | 317 | 317 | 92% | 4E-106 | 47.79 | 363 | NP_001030990.1 |
| unnamed protein product [Arabidopsis thaliana] | Arabidopsis thaliana | 316 | 316 | 92% | 8E-106 | 47.79 | 363 | CAD5312051.1 |
| mitogen-activated protein kinase homolog 2 [Arabidopsis thaliana] | Arabidopsis thaliana | 316 | 316 | 85% | 2E-105 | 49.68 | 376 | NP_564746.1 |
| ATMPK1 [Arabidopsis thaliana] | Arabidopsis thaliana | 315 | 315 | 93% | 2E-105 | 46.33 | 370 | CAA03535.1 |
| unnamed protein product [Arabidopsis thaliana] | Arabidopsis thaliana | 315 | 315 | 93% | 3E-105 | 46.33 | 370 | CAA0186271.1 |
| AT1G10210 [Arabidopsis thaliana] | Arabidopsis thaliana | 315 | 315 | 93% | 3E-105 | 46.33 | 370 | BAH19911.1 |
| mitogen-activated protein kinase 1 [Arabidopsis thaliana] | Arabidopsis thaliana | 315 | 315 | 93% | 3E-105 | 46.33 | 370 | NP_001031017.1 |
| mitogen-activated protein kinase 14 [Arabidopsis thaliana] | Arabidopsis thaliana | 315 | 315 | 95% | 3E-105 | 45.04 | 361 | NP_195363.1 |
| ATMPK2 [Arabidopsis thaliana] | Arabidopsis thaliana | 315 | 315 | 85% | 4E-105 | 49.35 | 376 | BAA03536.1 |
| unnamed protein product [Arabidopsis thaliana] | Arabidopsis thaliana | 313 | 313 | 95% | 1E-104 | 45.04 | 361 | CAA0397686.1 |
| MPK14 [Arabidopsis thaliana] | Arabidopsis thaliana | 311 | 311 | 95% | 8E-104 | 44.76 | 361 | OAO98488.1 |
| MAP kinase 5-CT [Arabidopsis thaliana] | Arabidopsis thaliana | 310 | 310 | 81% | 9E-104 | 50 | 337 | ADZ76491.1 |
| MAP kinase 7 [Arabidopsis thaliana] | Arabidopsis thaliana | 310 | 310 | 85% | 2E-103 | 48.71 | 368 | NP_179409.1 |
| MPK7 [Arabidopsis thaliana] | Arabidopsis thaliana | 310 | 310 | 85% | 2E-103 | 48.71 | 368 | OAP11028.1 |
| MAP Kinase [Arabidopsis thaliana] | Arabidopsis thaliana | 309 | 309 | 86% | 4E-103 | 50.48 | 354 | AAB61033.1 |
| unnamed protein product [Arabidopsis thaliana] | Arabidopsis thaliana | 310 | 310 | 91% | 4E-103 | 48.49 | 389 | CAD5321516.1 |
| unnamed protein product [Arabidopsis thaliana] | Arabidopsis thaliana | 309 | 309 | 85% | 5E-103 | 48.71 | 368 | CAA0364931.1 |
| MAP kinase 11 [Arabidopsis thaliana] | Arabidopsis thaliana | 307 | 307 | 78% | 2E-102 | 52.08 | 346 | NP_001322112.1 |
| MAP kinase [Arabidopsis thaliana] | Arabidopsis thaliana | 306 | 306 | 85% | 1E-101 | 48.06 | 368 | BAA04870.1 |
| unnamed protein product [Arabidopsis thaliana] | Arabidopsis thaliana | 310 | 310 | 86% | 3E-99 | 48.12 | 716 | CAD5330118.1 |
| mitogen-activated protein kinase 12 [Arabidopsis thaliana] | Arabidopsis thaliana | 297 | 297 | 80% | 5E-99 | 51.03 | 324 | NP_001189755.1 |
| mitogen-activated protein kinase 16 [Arabidopsis thaliana] | Arabidopsis thaliana | 287 | 287 | 91% | 7E-92 | 42.77 | 567 | NP_197402.1 |
| Unknown protein [Arabidopsis thaliana] | Arabidopsis thaliana | 287 | 287 | 91% | 7E-92 | 42.77 | 567 | AAL32607.1 |
| unnamed protein product [Arabidopsis thaliana] | Arabidopsis thaliana | 287 | 287 | 91% | 9E-92 | 42.77 | 569 | CAD5332107.1 |
| MAP kinase [Arabidopsis thaliana] | Arabidopsis thaliana | 276 | 276 | 75% | 6E-90 | 50 | 360 | BAD 95376.10 |
| putative MAP kinase [Arabidopsis thaliana] | Arabidopsis thaliana | 271 | 271 | 90% | 4E-87 | 42.6 | 431 | AAK62464.1 |
| MPK15 [Arabidopsis thaliana] | Arabidopsis thaliana | 274 | 274 | 92% | 1E-86 | 41.86 | 575 | OAP17567.1 |
| MAP kinase 19 [Arabidopsis thaliana] | Arabidopsis thaliana | 268 | 268 | 91% | 3E-86 | 42.18 | 425 | NP_001326968.1 |
| MAP kinase 15 [Arabidopsis thaliana] | Arabidopsis thaliana | 272 | 272 | 92% | 4E-86 | 41.86 | 576 | NP_565070.2 |
| MPK8 [Arabidopsis thaliana] | Arabidopsis thaliana | 269 | 269 | 92% | 1E-84 | 42.15 | 588 | OAP18797.1 |

## KEY RESOURCES TABLE

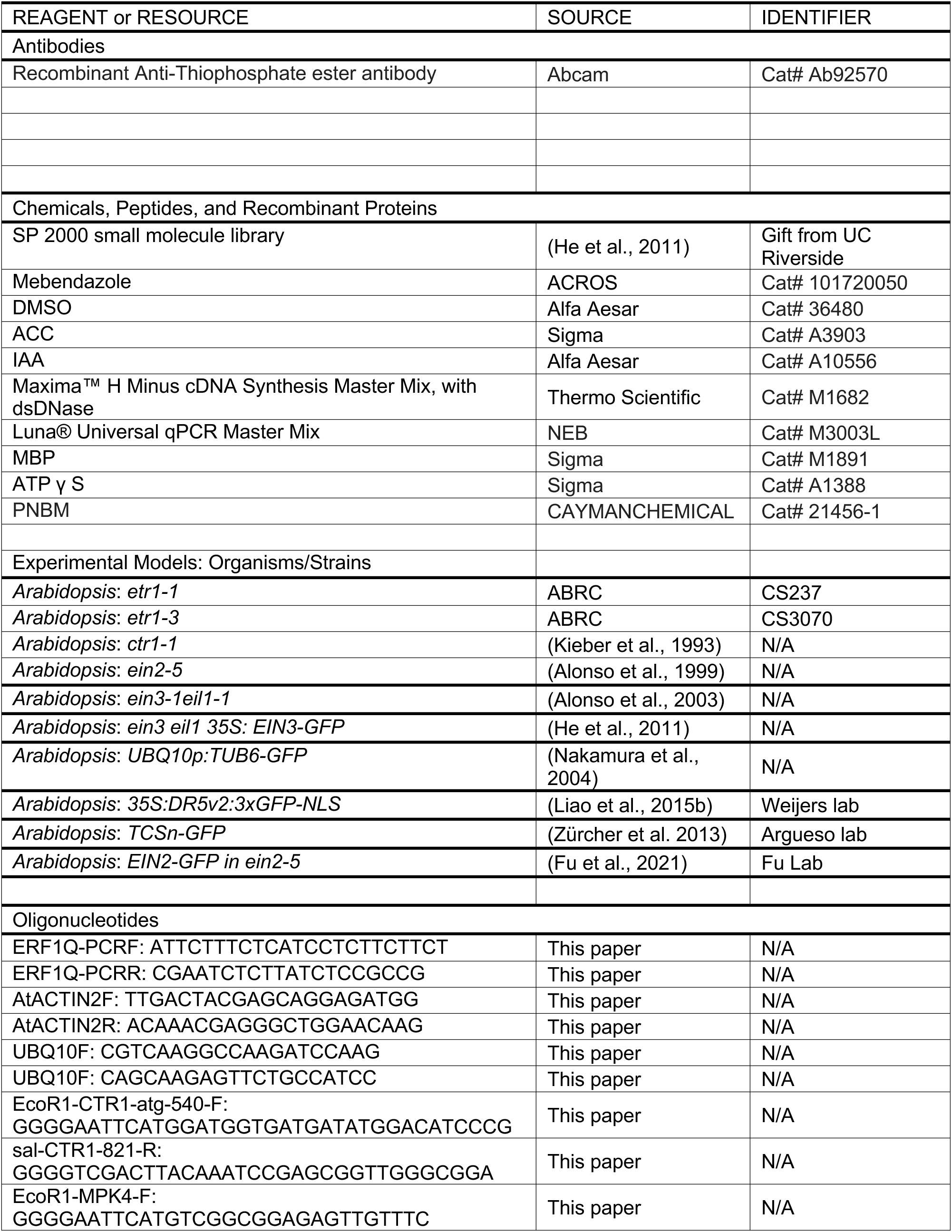
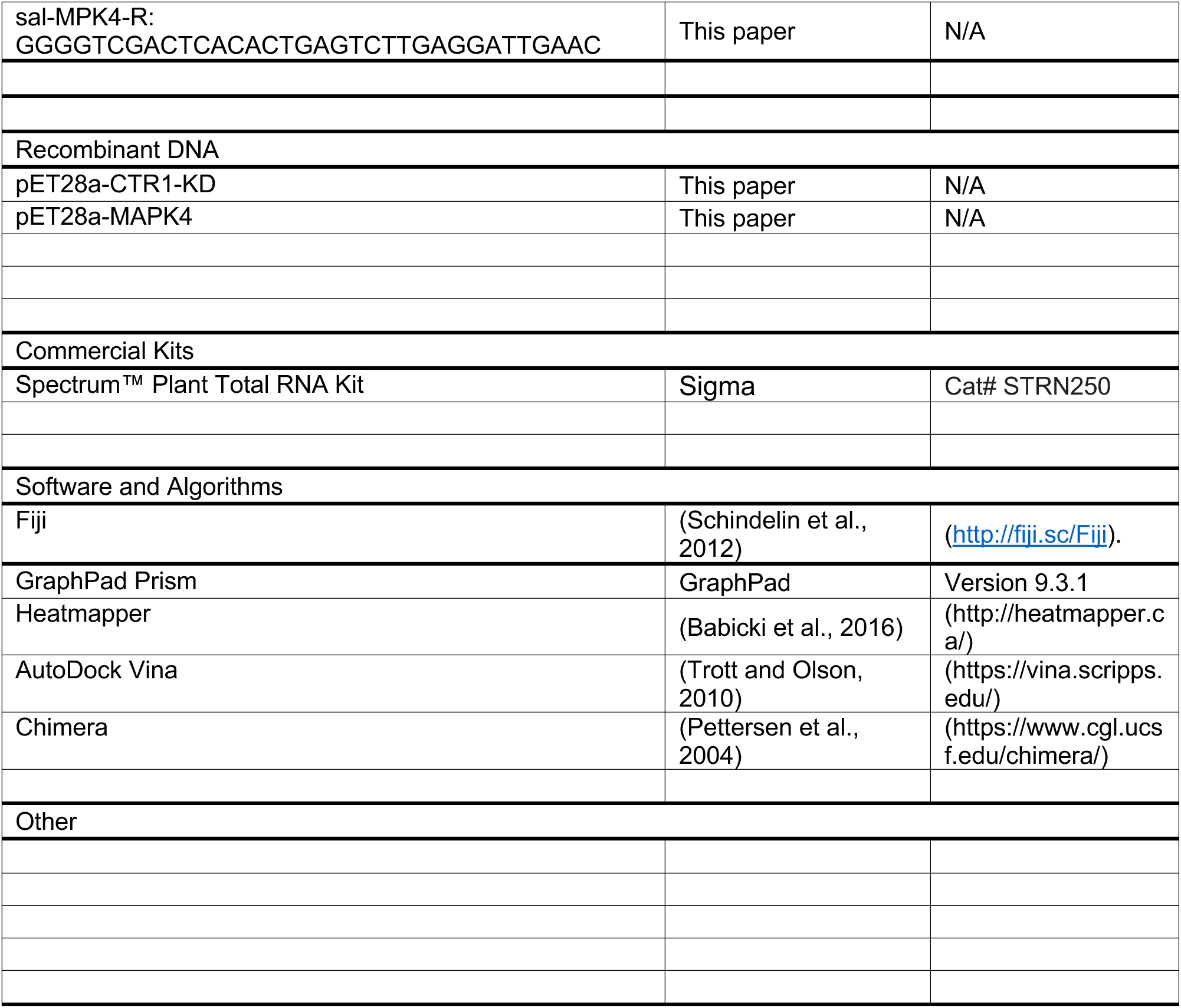

## CONTACT FOR REAGENT AND RESOURCE SHARING

Further information and requests for resources and reagents should be directed to and will be fulfilled by the Lead Contact, Wolfgang Busch.

## EXPERIMENTAL MODEL AND SUBJECT DETAILS

### Arabidopsis thaliana

Arabidopsis mutants and transgenic lines used in this study are in the Col-0 background if not mentioned otherwise. *etr1-1* (CS237) and *etr1-3* (CS3070) were ordered from ABRC, *ctr1-1* (Kieber et al., 1993) *ein2-5* (Alonso et al., 1999), *ein3-1 eil1-1*(Alonso et al., 2003), *ein3 eil1 35S: EIN3-GFP* (He et al., 2011), and *UBQ10p:TUB6-GFP* (Nakamura et al., 2004) were lab stock. *35S:DR5v2:3xGFP-NLS* (Liao et al., 2015b) was gift from Weijers lab (Col-Ultrecht background). *TCSn-GFP* (Zurcher et al., 2013) was a gift from Argueso lab*. EIN2-GFP* (Fu et al., 2021) in *ein2-5* was a gift from Fu lab. Plants were sown on half Murashige and Skoog (MS) medium with 1% (wt/vol) sugar, 0.1 % (wt/vol) MES (pH 5.75), containing 1% (wt/vol) agar supplemented with different chemicals as descripted in the article. Following a 3-day stratification in the cold room (4∼10°C), for green seedling assay, plants were grown on vertical plates in a walk-in growth chamber with long day conditions (16/8 h) at 21°C, 50 µM light intensity, 60% humidity. During nighttime, temperature was decreased to 15°C. For etiolated seedling assay, the plates were placed in light for 4 h and then germinated in the dark for 3 day in the growth chamber at 22 °C.

Plants were grown on vertical plates in a walk-in growth chamber with long day conditions (16/8 h) at 21°C, 50 µM light intensity, 60% humidity. During nighttime, temperature was decreased to 15°C.

### Accession Numbers

Sequence data from this article can be found in the Arabidopsis Genome Initiative or GenBank/EMBL databases or Uniprot databases under the following accession numbers: *ERF1* (AT3G23240), *ETR1* (AT1G66340), *CTR1* (AT5g03730), *EIN2* (AT5G03280), *EIN3* (AT3g20770), *EIL1* (AT2G27050), *MAPK4* (AT4G01370), and *hsMAPK14* (HGNC:6876).

Accession numbers of protein and chemicals structures used in the molecular modeling in Figure 6 are as follows: CTR1 (PDB: 3PPZ), MAPK14 (PDB: 6SFO) and MBZ (CHEMBL685).

## METHOD DETAILS

### Small Molecule Library and Screen Information

The SP2000 small molecule library was received from the University of California, Riverside. The library was first screened at a concentration of 50 ∼ 100 µM in 200 µL of liquid MS media using wild type Arabidopsis. After being grown in the liquid media with small molecules for 7 days, seedlings were transferred to vertical plates to grow for another 2 days. The root phenotypes were acquired by CCD flatbed scanners. After candidates were picked, a second round of screen was performed by growing Col-0 seedlings on vertical plates with gradient concentration of candidate small molecules, from 1 µM to 100 µM, for 14 days to confirm the phenotype and optimal concentration.

### Cloning and constructs

To produce recombinant CTR1 kinase domain (CTR1-KD) and MAPK4 proteins in *E. coli*, the coding sequences of 540-821 aa of CTR1and full length MAPK4 were PCR amplified from cDNA of Arabidopsis Col-0, using gene-specific primers (listed in KEY RESOURCES TABLE). The PCR products were inserted into pET28a using restriction digestion and ligation with EcoR 1 and Sal 1 (NEB) sites.

### Chemical Solutions

The stock solutions for preparing media for the plant experiments were Mebendazole (10 mM) in DMSO, ACC (10 mM) in dH2O, IAA (1 mM) in ethanol.

### Root phenotyping

Images of root phenotypes on plates were acquired with CCD flatbed scanners (EPSON). Root lengths and lateral root angles were measured using Fiji (http://fiji.sc/Fiji).

### Lateral root angle measurement

The lateral root angle of individual LRs was measured according to (Rosquete et al., 2013b) using Fiji. LRs after stage III from each seedling were measured according to (Rosquete et al., 2013b; Waidmann et al., 2019a) .

### Time lapse of lateral root angle phenotyping

14-day-old seedlings on DMSO plate or 19 d old seedling on mebendazole plate were transferred to DMSO or 1 µM mebendazole plates as described (Figure 1E). Root phenotype images were acquired by CCD vertical scanners (EPSON) every 10 mins over a period of 24 hours using the tool described(Ogura et al., 2022). Root length and lateral root angles were measured using Fiji (http://fiji.sc/Fiji).

### mRNA-seq

#### A#Tissue preparation

Roots or lateral root tips of 14-day-old Col-0 seedlings with different treatment were collected for the mRNA-seq analysis. In detail:

##### a#Roots

Sterilized seeds of Col-0 on ½ MS media were planted on a mesh in one row (∼more than 30 seeds/plate). 5 plates were used for one treatment condition for every replicate, and 4 replicates were used for every treatment. After 14 days of vertical growth in the chamber, liquid media of ½ MS with DMSO (0.1 %, v/v), 10 µM MBZ, and 10 µM ACC (to be consistent, we also added the same concentration of DMSO (0.1 %, v/v) in the ACC treatment plates) were prepared. 10 mL liquid treatment medium was used on each plate. The mesh with seedlings from ½ MS plates was transferred the respective liquid treatment plates and kept in growth chamber. After 4 hours for the mRNA-seq of MBZ treatment and 3 hours for the mRNA-seq of MBZ and ACC treatment, roots were collected using scalpel and tweezers. Excess liquid on the roots was removed using a paper towel. Root samples were flash-frozen in liquid nitrogen and then stored at -80 freezer before RNA extraction.

For RNA extraction, root tissues were ground in liquid nitrogen and RNA was extracted using Quick-RNA Kits (Zymo research). RNA libraries were prepared for sequencing using standard Illumina protocols at the Salk Next Generation Sequencing Core (NGS). Note: We used liquid media for DMSO, MBZ, and ACC treatment in the mRNA-seq of roots to detect transient gene regulation as these chemicals in liquid media can be sufficiently and equally absorbed into lateral roots and reduce time for sample collection.

##### b#Lateral root tips

Sterilized seeds were grown for 14 days on ½ MS medium. The seedlings were then transferred to freshly prepared DMSO or 1µM MBZ plates for 4 hours on 15 plates (5 plants/plate) for each treatment. 4 biological replications were performed. A stereo microscope (Leica) was used to dissect the meristem area of lateral root tips in order to collect the lateral root tips. The cut lateral root tips were quickly placed into 1.5 microcentrifuge tubes that floated on liquid nitrogen. This process was repeated until approximately 400 root tips were collected per sample.

The NucleoSpin RNA extraction kit (Macherey-Nagel) was used to extract total RNA. The concentration and integrity of extracted RNA were determined using Qubit and TapeStation 2200 before being sent to Salk Next Generation Sequencing Core (NGS) for library preparation and sequencing.

#### B#Data analysis

##### a#Read alignment

The TAIR10 genome file and annotation file were obtained from the Arabidopsis Information resource web site (http://www.arabidopsis.org)(Berardini et al., 2015). The An aligner, called the Splice Transcripts Alignments to Reference (STAR) version 2.7.0a (Dobin et al., 2013), was used to align short reads in the FASTQ files.

##### b#Differential expression analysis

Differential expression of genes between treatments and control was determined using the R package, *edgeR* (version 3.36.0)(Robinson et al., 2010). The *CPM* (counts per million) function from *edgeR* was used to normalize the counts and differentially expressed genes (DEGs) were identified using *glmLRT* function from edgeR. A false discovery rate (FDR < 0.05) and |log2FC| > 1 were used as the criterial values for identification of up-regulated and down-regulated DEGs.

### GO analysis

Gene Ontology analysis of up-regulated DEGs (FDR < 0.05) and |log2FC| > 1 by MBZ treatment in roots of 14-day-old Arabidopsis seedlings was performed with PANTHER Overrepresentation Test (Released 20230705). The Annotation Version used GO Ontology database (DOI: 10.5281/zenodo.7942786 (Released 20230510)). Enriched genes (Bonferroni-corrected for P < 0.05) for biological processes were listed (Figure 2A). We used R package topGO (version 2.42.0)(Alexa et al., 2006) with the weight01 algorithm and the mappings from the “org.At.tair.db” R package to perform functional enrichment analysis of DEGs by MBZ treatment in roots and lateral root tips of 14-day-old Arabidopsis seedlings. We carried out KS test for statistical comparison. Then we used “ggplot2” package in R to plot 30 most enriched gene ontology terms (GO) which were ordered by Kolmogorov-Smirnov p-values for biological processes (Figure 2F, S2D).

### Measurement of ethylene production

100 seeds of Col-0 were planted and sealed in a screw-top 20 mL vial containing 10 mL ½ MS medium supplemented with DMSO (control), 10 µM mebendazole, and 10 µM ACC treatments (4 replicates). These vials were incubated at 22 °C for 3 days under dark environment. 500-1000 µL sample of the headspace gas was sampled with an autosampler (TriPlus RSH, Thermo Scientific) and injected into a gas chromatograph (Trace 1310 GC, Thermo Scientific) that was equipped with a HP-PLOTQ column (30 mm, 320 µm, 20 µm) and a mass spectrometer (TSQ8000 Evo MS, Thermo Scientific), scanning from 25-27.5 m/z. Separations were carried out at 35°C using He as the carrier gas. The area of the ethylene peak (RT: 4 min) was integrated using Thermo Xcalibur Qual Browser. A calibration curve was generated by varying the injection volume (100 µL, 250 µL, 500 µL, and 1000 µL) of a 10 ppm ethylene standard (in nitrogen), and sample results are expressed as concentrations calculated from linear regression of calibration samples.

### qRT-PCR

For measurement of marker genes of plant hormone pathways, 20 7d old seedlings of Col-0 grown on ½ MS media were treated with DMSO or 10µM MBZ, dH2O or 10 µM ACC for 2h, or 3, 6, 12h as described in the manuscript. Then whole roots were collected. RNA extraction was performed using the Spectrum™ Plant Total RNA Kit (Sigma), followed by cDNA synthesis using Maxima™ H Minus cDNA Synthesis Master Mix, with dsDNase (Thermo Scientific). Quantitative PCR was performed with Luna® Universal qPCR Master Mix (NEB). All procedures were carried out according to manufacturer instructions. qRT-PCR primer information were listed in the KEY RESOURCES TABLE.

### Microscopy

A Zeiss 710 confocal microscope with a 40x objective was used to detect propidium iodide (PI) fluorescence in Figure 3F. A Keyence BZ-X810 microscope with a 20x objective was used to detect bright field images and GFP fluorescence in Figure 3B, Figure S1B right panel. A Leica TCS SP5 confocal microscope with 40x objective was used to detect GFP fluorescence in Figure 7A.

### Accumulation of EIN2-GFP in root cells

3-day-old etiolated seedlings of two independent transgenic *EIN2-GFP* reporter lines in which the GFP was fused to EIN2’s C-terminus (*pEIN2::EIN2-GFP* in the *ein2-5* background (Fu et al., 2021)) were moved into different wells of 6-well plate with 5 mL Mock (dH2O+ 5 µL DMSO), ACC (100 µM + 5 µL DMSO), or MBZ (10 µM). The plate was then covered with tin foil to keep the dark environment for 3 hour. Then samples were directly mounted on glass slides in their treatment liquid and observed under a Leica SP5 microscope (Leica).

### Time lapse microscopy of reporter genes in lateral root tips

To observe the response of ethylene (*EIN3-GFP*), auxin (*DR5v2-GFP*), and cytokinin (*TCSn-GFP*) reporter lines to MBZ, 14-day-old seedlings grown on ½ MS media were transferred to a Chambered Coverglass (Thermo Scientific) that was filled with treatment medium. For chamber preparation, 6 mL of ½ MS medium (1 % agar) supplemented with the different treatments (DMSO, 10 µM MBZ, 1 µM IAA, 50 µM ACC, and 10 µM tZ) was filled to each chamber. After 20 minutes of medium solidification, the seedlings were placed on the media surface and the lateral roots were adjusted to positions suitable for imaging.

For imaging, a Zeiss CSU Spinning Disk Confocal Microscope equipped with laser 488 mm and a Yokogawa spinning disk scan head with EM-CCD camera were used to set up the image acquisition every 20 minutes for a total of 4 hours. Three Z-stack positions were automatically captured at each time point under 10X objective and Fiji was used to quantify the average intensity of GFP fluorescence. For proper comparison at different treatment conditions of each reporter lines, we normalized the average fluorescence data as the percentages (or relative fluorescence intensity as text label). In detail, the first time point of each treatment is set as 100% and the relative fluorescence intensity from the second time point is calculated as the fold-change in percentages comparing the first time point.

### Quantification of relative GFP fluorescent intensity between the nucleus and cytosol

The quantification of the microscopy localization data of *EIN2-GFP* in Figure 7B was done according to (Fu et al., 2021) using LAS X (Leica Application Suite v. X3.1.1.15751) and GraphPad Prism V9.5.1. In brief, the signal intensity of nucleus (Ni) and whole cell (Wi) was quantified by LAS X (Mean value of nucleus * ROI Area of nucleus, mean value of whole cell * ROI Area of whole cell), and then the signal intensity of cytosol (Ci) was obtained by Wi minus Ni, and the area of cytosol (Sc) was obtained by the area of whole cell (ROI Area of whole cell) minus the area of nucleus (ROI Area of nucleus). The mean value of the signal intensity of cytosol (Mean Ci) equals Ci/Sc, and the relative GFP fluorescent intensity between the nucleus and cytosol equals Mean value of nucleus/ Mean Ci.

### Computational Docking and Molecular Modeling

The AutoDock Vina and Chimera software were used to dock mebendazole (CHEMBL685) into crystal structure of CTR1 (PDB: 3PPZ) and MAPK14 (PDB: 6SFO). Figures for those structures were visualized and generated by Chimera and Pymol.

### Protein purification

The CTR1 Kinase domain (540 aa-821 aa) (CTR1-KD) and MAPK4 were cloned into a pET28a. Proteins were expressed in Rosetta™ 2(DE3)pLysS Competent Cells. Cultures were incubated at 37°C (OD 600:0.7 to 0.9) and induced by adding 0.4 mM isopropylthioβ-D-galactoside (IPTG) at 20°C for 22 h, 200 rpm. Cells were harvested and resuspended in lysis buffer (50 mM pH 7.5 Tris-HCl, 1mM EDTA, 100 mM NaCl, 20% glycerol, EDTA-free protease inhibitor cocktail, 10 mM imidazole). followed by sonication and centrifugation (13,000 × g × 20 min). The His-tagged proteins were first purified using QIAGEN Ni-NTA Agarose. The elute from Ni-NTA was collected and further purified by FPLC using superdex 200 Increase 10/300 column (Mayerhofer et al., 2011). The fractions corresponding to single peak from FPLC were combined, and concentrated to a desired volume. Protein purity was verified by Coomassie blue gel staining. Protein concentration was measured by their absorbance at 280 nm.

### *In vitro* kinase assay with non-radioactive ATP-γ-S

The *in vitro* kinase assay followed the previous published method (Allen et al., 2007). 100 ng purified recombinant CTR1-KD or 1 µg MPK4 was mixed with 2 µg Myelin Basic Protein (MBP, a common kinase substrate, Sigma Cat. M1891) and 1 mM ATPγS in kinase reaction buffer (10 mM HEPES pH 7.4, 150 mM NaCl, 10 mM MgCl2, 1xRoche Complete Protease Inhibitor mixture) and different concentration of MBZ at room temperature and vortexed for 1 hour. The reactions were terminated by 20 mM EDTA, and the proteins were alkylated by 1.5 µL of 50 mM PNBM (CAYMAN CHEMICAL CO, Cat. 21456-1) at room temperature and vortexed for 2 hours. This step was terminated by denaturing in NuPAGE™ LDS Sample Buffer and 1x NuPAGE™ Sample Reducing Agent (Invitrogen™). Samples were subjected to SDS-PAGE. The phosphorylation status of MBP, CTR1-KD and MPK4 were analyzed by western blot using Recombinant Anti-Thiophosphate ester antibody (Abcam, Cat. Ab92570, 1:5000).

### Statistical analysis

A two-sided Student’s t-test was used for comparisons of two conditions. The comparison of meristem length, mature cell length, root length, GSA between multiple treatments or genotypes was performed by One-way ANOVA and *post hoc* Tukey testing. Detailed information was prepared in figure legends. Statistical significance of overlap between DEGs from MBZ treatment and ethylene treatment was assessed by hypergeometric distribution test.

## Data availability

Data that support the findings of this study have been deposited in GEO. The GEO number is GSE201648 .

## References

Alexa, A., Rahnenfuhrer, J., and Lengauer, T. (2006). Improved scoring of functional groups from gene expression data by decorrelating GO graph structure. Bioinformatics 22, 1600–1607.

Allen, J.J., Li, M., Brinkworth, C.S., Paulson, J.L., Wang, D., Hubner, A., Chou, W.H., Davis, R.J., Burlingame, A.L., Messing, R.O., et al. (2007). A semisynthetic epitope for kinase substrates. Nat Methods 4, 511–516.

Alonso, J.M., Hirayama, T., Roman, G., Nourizadeh, S., and Ecker, J.R. (1999). EIN2, a bifunctional transducer of ethylene and stress responses in Arabidopsis. Science 284, 2148–2152.

Alonso, J.M., Stepanova, A.N., Solano, R., Wisman, E., Ferrari, S., Ausubel, F.M., and Ecker, J.R. (2003). Five components of the ethylene-response pathway identified in a screen for weak ethylene-insensitive mutants in Arabidopsis. Proc Natl Acad Sci U S A 100, 2992–2997.

An, F., Zhao, Q., Ji, Y., Li, W., Jiang, Z., Yu, X., Zhang, C., Han, Y., He, W., Liu, Y., et al. (2010). Ethylene-induced stabilization of ETHYLENE INSENSITIVE3 and EIN3- LIKE1 is mediated by proteasomal degradation of EIN3 binding F-box 1 and 2 that requires EIN2 in Arabidopsis. Plant Cell 22, 2384–2401.

Ariey-Bonnet, J., Carrasco, K., Le Grand, M., Hoffer, L., Betzi, S., Feracci, M., Tsvetkov, P., Devred, F., Collette, Y., Morelli, X., et al. (2020). In silico molecular target prediction unveils mebendazole as a potent MAPK14 inhibitor. Mol Oncol 14, 3083–3099.

Babicki, S., Arndt, D., Marcu, A., Liang, Y.J., Grant, J.R., Maciejewski, A., and Wishart, D.S. (2016). Heatmapper: web-enabled heat mapping for all. Nucleic Acids Res 44, W147–W153.

Berardini, T.Z., Reiser, L., Li, D.H., Mezheritsky, Y., Muller, R., Strait, E., and Huala, E. (2015). The arabidopsis information resource: Making and mining the “gold standard” annotated reference plant genome. Genesis 53, 474–485.

Chang, C., Kwok, S.F., Bleecker, A.B., and Meyerowitz, E.M. (1993). Arabidopsis ethylene-response gene ETR1: similarity of product to two-component regulators. Science 262, 539–544.

Chang, K.N., Zhong, S., Weirauch, M.T., Hon, G., Pelizzola, M., Li, H., Huang, S.S.C., Schmitz, R.J., Urich, M.A., Kuo, D., et al. (2013). Temporal transcriptional response to ethylene gas drives growth hormone cross-regulation in Arabidopsis. Elife 2.

Chao, Q., Rothenberg, M., Solano, R., Roman, G., Terzaghi, W., and Ecker, J.R. (1997). Activation of the ethylene gas response pathway in Arabidopsis by the nuclear protein ETHYLENE-INSENSITIVE3 and related proteins. Cell 89, 1133–1144.

De Cnodder, T., Vissenberg, K., Van Der Straeten, D., and Verbelen, J.P. (2005). Regulation of cell length in the Arabidopsis thaliana root by the ethylene precursor 1- aminocyclopropane- 1-carboxylic acid: a matter of apoplastic reactions. The New phytologist 168, 541–550.

Dobin, A., Davis, C.A., Schlesinger, F., Drenkow, J., Zaleski, C., Jha, S., Batut, P., Chaisson, M., and Gingeras, T.R. (2013). STAR: ultrafast universal RNA-seq aligner. Bioinformatics 29, 15–21.

Feng, Y., Xu, P., Li, B., Li, P., Wen, X., An, F., Gong, Y., Xin, Y., Zhu, Z., Wang, Y., et al. (2017). Ethylene promotes root hair growth through coordinated EIN3/EIL1 and RHD6/RSL1 activity in Arabidopsis. Proc Natl Acad Sci U S A 114, 13834–13839.

Fu, L.W., Liu, Y.L., Qin, G.C., Wu, P., Zi, H.L., Xu, Z.T., Zhao, X.D., Wang, Y., Li, Y.X., Yang, S.H., et al. (2021). The TOR-EIN2 axis mediates nuclear signalling to modulate plant growth. Nature 591, 288–292.

Furutani, M., and Morita, M.T. (2021). LAZY1-LIKE-mediated gravity signaling pathway in root gravitropic set-point angle control. Plant Physiology 187, 1087–1095.

Ge, L.F., and Chen, R.J. (2016). Negative gravitropism in plant roots. Nature plants 2.

Guo, H., and Ecker, J.R. (2003). Plant responses to ethylene gas are mediated by SCF(EBF1/EBF2)-dependent proteolysis of EIN3 transcription factor. Cell 115, 667–677.

Guseman, J.M., Webb, K., Srinivasan, C., and Dardick, C. (2017). DRO1 influences root system architecture in Arabidopsis and Prunus species. Plant J 89, 1093–1105.

Guzman, P., and Ecker, J.R. (1990). Exploiting the triple response of Arabidopsis to identify ethylene-related mutants. Plant Cell 2, 513–523.

He, W.R., Brumos, J., Li, H.J., Ji, Y.S., Ke, M., Gong, X.Q., Zeng, Q.L., Li, W.Y., Zhang, X.Y., An, F.Y., et al. (2011). A Small-Molecule Screen Identifies L-Kynurenine as a Competitive Inhibitor of TAA1/TAR Activity in Ethylene-Directed Auxin Biosynthesis and Root Growth in Arabidopsis. Plant Cell 23, 3944–3960.

Hua, J., Sakai, H., Nourizadeh, S., Chen, Q.G., Bleecker, A.B., Ecker, J.R., and Meyerowitz, E.M. (1998). EIN4 and ERS2 are members of the putative ethylene receptor gene family in Arabidopsis. Plant Cell 10, 1321–1332.

Ju, C., Van de Poel, B., Cooper, E.D., Thierer, J.H., Gibbons, T.R., Delwiche, C.F., and Chang, C. (2015). Conservation of ethylene as a plant hormone over 450 million years of evolution. Nature plants 1, 14004.

Ju, C., Yoon, G.M., Shemansky, J.M., Lin, D.Y., Ying, Z.I., Chang, J., Garrett, W.M., Kessenbrock, M., Groth, G., Tucker, M.L., et al. (2012). CTR1 phosphorylates the central regulator EIN2 to control ethylene hormone signaling from the ER membrane to the nucleus in Arabidopsis. Proc Natl Acad Sci U S A 109, 19486–19491.

Kieber, J.J., Rothenberg, M., Roman, G., Feldmann, K.A., and Ecker, J.R. (1993). CTR1, a negative regulator of the ethylene response pathway in Arabidopsis, encodes a member of the raf family of protein kinases. Cell 72, 427–441.

Le, J., Vandenbussche, F., Van Der Straeten, D., and Verbelen, J.P. (2001). In the early response of Arabidopsis roots to ethylene, cell elongation is up- and down-regulated and uncoupled from differentiation. Plant Physiol 125, 519–522.

Li, W., Lacey, R.F., Ye, Y., Lu, J., Yeh, K.C., Xiao, Y., Li, L., Wen, C.K., Binder, B.M., and Zhao, Y. (2017). Triplin, a small molecule, reveals copper ion transport in ethylene signaling from ATX1 to RAN1. PLoS Genet 13, e1006703.

Liao, C.Y., Smet, W., Brunoud, G., Yoshida, S., Vernoux, T., and Weijers, D. (2015a). Reporters for sensitive and quantitative measurement of auxin response. Nat Methods 12, 207–210, 202 p following 210.

Liao, C.Y., Smet, W., Brunoud, G., Yoshida, S., Vernoux, T., and Weijers, D. (2015b). Reporters for sensitive and quantitative measurement of auxin response (vol 12, pg 207, 2015). Nat Methods 12, 1098–1098.

Liao, C.Y., Smet, W., Brunoud, G., Yoshida, S., Vernoux, T., and Weijers, D. (2015). Reporters for sensitive and quantitative measurement of auxin response. Nat Methods 12, 207–210, 202 p following 210.

Lynch, J.P. (2019). Root phenotypes for improved nutrient capture: an underexploited opportunity for global agriculture. New Phytol 223, 548–564.

Mayerhofer, H., Mueller-Dieckmann, C., and Mueller-Dieckmann, J. (2011). Cloning, expression, purification and preliminary X-ray analysis of the protein kinase domain of constitutive triple response 1 (CTR1) from Arabidopsis thaliana. Acta Crystallogr Sect F Struct Biol Cryst Commun 67, 117–120.

Mayerhofer, H., Panneerselvam, S., and Mueller-Dieckmann, J. (2012). Protein kinase domain of CTR1 from Arabidopsis thaliana promotes ethylene receptor cross talk. J Mol Biol 415, 768–779.

Mou, W., Kao, Y.T., Michard, E., Simon, A.A., Li, D., Wudick, M.M., Lizzio, M.A., Feijo, J.A., and Chang, C. (2020). Ethylene-independent signaling by the ethylene precursor ACC in Arabidopsis ovular pollen tube attraction. Nat Commun 11, 4082.

Muller, M., and Munne-Bosch, S. (2015). Ethylene Response Factors: A Key Regulatory Hub in Hormone and Stress Signaling. Plant Physiol 169, 32–41.

Nakamura, M., Naoi, K., Shoji, T., and Hashimoto, T. (2004). Low concentrations of propyzamide and oryzalin alter microtubule dynamics in Arabidopsis epidermal cells. Plant Cell Physiol 45, 1330–1334.

Negi, S., Ivanchenko, M.G., and Muday, G.K. (2008). Ethylene regulates lateral root formation and auxin transport in Arabidopsis thaliana. Plant J 55, 175–187.

Ogura, T., Goeschl, C., and Busch, W. (2022). A Multiplexed, Time-Resolved Assay of Root Gravitropic Bending on Agar Plates. Methods Mol Biol 2368, 61–70.

Pattyn, J., Vaughan-Hirsch, J., and Van de Poel, B. (2021). The regulation of ethylene biosynthesis: a complex multilevel control circuitry. New Phytol 229, 770–782.

Pettersen, E.F., Goddard, T.D., Huang, C.C., Couch, G.S., Greenblatt, D.M., Meng, E.C., and Ferrin, T.E. (2004). UCSF Chimera--a visualization system for exploratory research and analysis. J Comput Chem 25, 1605–1612.

Poirier, V., Roumet, C., and Munson, A.D. (2018). The root of the matter: Linking root traits and soil organic matter stabilization processes. Soil Biology and Biochemistry 120, 246–259.

Qiao, H., Shen, Z., Huang, S.S., Schmitz, R.J., Urich, M.A., Briggs, S.P., and Ecker, J.R. (2012a). Processing and subcellular trafficking of ER-tethered EIN2 control response to ethylene gas. Science 338, 390–393.

Qiao, H., Shen, Z.X., Huang, S.S.C., Schmitz, R.J., Urich, M.A., Briggs, S.P., and Ecker, J.R. (2012b). Processing and Subcellular Trafficking of ER-Tethered EIN2 Control Response to Ethylene Gas. Science 338, 390–393.

Robinson, M.D., McCarthy, D.J., and Smyth, G.K. (2010). edgeR: a Bioconductor package for differential expression analysis of digital gene expression data. Bioinformatics 26, 139–140.

Rosquete, M.R., von Wangenheim, D., Marhavy, P., Barbez, E., Stelzer, E.H., Benkova, E., Maizel, A., and Kleine-Vehn, J. (2013a). An auxin transport mechanism restricts positive orthogravitropism in lateral roots. Current biology : CB 23, 817–822.

Rosquete, M.R., von Wangenheim, D., Marhavy, P., Barbez, E., Stelzer, E.H.K., Benkova, E., Maizel, A., and Kleine-Vehn, J. (2013b). An Auxin Transport Mechanism Restricts Positive Orthogravitropism in Lateral Roots. Curr Biol 23, 817–822.

Rosquete, M.R., von Wangenheim, D., Marhavy, P., Barbez, E., Stelzer, E.H., Benkova, E., Maizel, A., and Kleine-Vehn, J. (2013). An auxin transport mechanism restricts positive orthogravitropism in lateral roots. Current biology : CB 23, 817–822.

Roychoudhry, S., Del Bianco, M., Kieffer, M., and Kepinski, S. (2013). Auxin controls gravitropic setpoint angle in higher plant lateral branches. Current biology : CB 23, 1497–1504.

S F Yang, a., and Hoffman, N.E. (1984). Ethylene Biosynthesis and its Regulation in Higher Plants. Annual Review of Plant Physiology 35, 155–189.

Sakai, H., Hua, J., Chen, Q.G., Chang, C., Medrano, L.J., Bleecker, A.B., and Meyerowitz, E.M. (1998). ETR2 is an ETR1-like gene involved in ethylene signaling in Arabidopsis. Proc Natl Acad Sci U S A 95, 5812–5817.

Schindelin, J., Arganda-Carreras, I., Frise, E., Kaynig, V., Longair, M., Pietzsch, T., Preibisch, S., Rueden, C., Saalfeld, S., Schmid, B., et al. (2012). Fiji: an open-source platform for biological-image analysis. Nat Methods 9, 676–682.

Sun, X.Z., Li, Y.X., He, W.R., Ji, C.G., Xia, P.X., Wang, Y.C., Du, S., Li, H.J., Raikhel, N., Xiao, J.Y., et al. (2017). Pyrazinamide and derivatives block ethylene biosynthesis by inhibiting ACC oxidase. Nat Commun 8.

Trott, O., and Olson, A.J. (2010). AutoDock Vina: improving the speed and accuracy of docking with a new scoring function, efficient optimization, and multithreading. J Comput Chem 31, 455–461.

Uga, Y., Sugimoto, K., Ogawa, S., Rane, J., Ishitani, M., Hara, N., Kitomi, Y., Inukai, Y., Ono, K., Kanno, N., et al. (2013). Control of root system architecture by DEEPER ROOTING 1 increases rice yield under drought conditions. Nature genetics 45, 1097–1102.

Vaseva, II, Qudeimat, E., Potuschak, T., Du, Y., Genschik, P., Vandenbussche, F., and Van Der Straeten, D. (2018). The plant hormone ethylene restricts Arabidopsis growth via the epidermis. Proc Natl Acad Sci U S A 115, E4130–E4139.

Waidmann, S., Rosquete, M.R., Scholler, M., Sarkel, E., Lindner, H., LaRue, T., Petrik, I., Dunser, K., Martopawiro, S., Sasidharan, R., et al. (2019a). Cytokinin functions as an asymmetric and anti-gravitropic signal in lateral roots. Nat Commun 10.

Waidmann, S., Ruiz Rosquete, M., Scholler, M., Sarkel, E., Lindner, H., LaRue, T., Petrik, I., Dunser, K., Martopawiro, S., Sasidharan, R., et al. (2019b). Cytokinin functions as an asymmetric and anti-gravitropic signal in lateral roots. Nat Commun 10, 3540.

Waidmann, S., Ruiz Rosquete, M., Scholler, M., Sarkel, E., Lindner, H., LaRue, T., Petrik, I., Dunser, K., Martopawiro, S., Sasidharan, R., et al. (2019). Cytokinin functions as an asymmetric and anti-gravitropic signal in lateral roots. Nat Commun 10, 3540.

Waite, J.M., and Dardick, C. (2021). The roles of the IGT gene family in plant architecture: past, present, and future. Curr Opin Plant Biol 59.

Wen, X., Zhang, C., Ji, Y., Zhao, Q., He, W., An, F., Jiang, L., and Guo, H. (2012). Activation of ethylene signaling is mediated by nuclear translocation of the cleaved EIN2 carboxyl terminus. Cell Res 22, 1613–1616.

Xiao, G., and Zhang, Y. (2020). Adaptive Growth: Shaping Auxin-Mediated Root System Architecture. Trends in plant science 25, 121–123.

Yoshihara, T., and Spalding, E.P. (2017). LAZY Genes Mediate the Effects of Gravity on Auxin Gradients and Plant Architecture. Plant Physiology 175, 959–969.

Zhu, Y., Li, H.J., Su, Q., Wen, J., Wang, Y.F., Song, W., Xie, Y.P., He, W.R., Yang, Z., Jiang, K., et al. (2019). A phenotype-directed chemical screen identifies ponalrestat as an inhibitor of the plant flavin monooxygenase YUCCA in auxin biosynthesis. J Biol Chem 294, 19923–19933.

Zurcher, E., Tavor-Deslex, D., Lituiev, D., Enkerli, K., Tarr, P.T., and Muller, B. (2013). A robust and sensitive synthetic sensor to monitor the transcriptional output of the cytokinin signaling network in planta. Plant Physiol 161, 1066–1075.

